# Harmonic representations of regions and interactions in spatial transcriptomics

**DOI:** 10.1101/2024.08.14.607982

**Authors:** Kamal Maher, Xiao Wang

## Abstract

Spatial transcriptomics technologies enable unbiased measurement of the cell-cell interactions underlying tissue structure and function. However, most unsupervised methods instead focus on identifying tissue regions, representing them as positively covarying low-frequency spatial patterns of gene expression over the tissue. Here, we extend this frequency-based (i.e. harmonic) approach to show that *negatively* covarying *high* frequencies represent interactions. Similarly, combinations of low and high frequencies represent interactions along large length scales, or, equivalently, region boundaries. The resulting equations further reveal a duality in which regions and interactions are complementary representations of the same underlying information, each with unique strengths and weaknesses. We demonstrate these concepts in multiple datasets from human lymph node, human tonsil, and mouse models of Alzheimer’s disease. Altogether, this work offers a conceptually consistent quantitative framework for spatial transcriptomics.

## Introduction

Cell-cell interactions determine tissue structure and function. They drive the very formation of tissues during development (Perrimon et al., 2012), shape the progression of disease (de Visser & Joyce, 2023), and enable coordinated immune responses (Schenkel & Pauken, 2023). Unbiased measurement of such interactions could enable hypothesis generation regarding the influence of interactions on tissue function. This is the promise of spatial transcriptomics technologies (Marx, 2021; Tian et al., 2023; Larsson et al., 2021; Palla et al., 2022a; Rao et al., 2021). However, unsupervised quantification of interactions in spatial transcriptomics data remains a challenge.

Few fully unsupervised methods exist for quantifying interactions. The majority rely on a single principle: calculating the spatial covariance between cell types (Palla et al., 2022b; Dries et al., 2021; Pham et al., 2023; Schapiro et al., 2017; Miller et al., 2021; Schürch et al., 2020; Ali et al., 2020). This amounts to counting the instances in which one cell type is next to another cell type in the tissue, yielding a type-by-type spatial covariance matrix. However, predetermined cell types obscure the rich gene-level information provided by transcriptomics technologies. One might wonder whether this approach could instead be generalized to the gene level, perhaps by calculating a gene-by-gene spatial covariance matrix (Chen, 2015; Russell et al., 2023). This matrix might then be distilled into interaction-specific gene programs via methods such as principal components analysis (PCA). However, there is a fundamental problem with this approach: the spatial covariance matrix is indefinite, causing PCA to produce imaginary numbers and negative variance explained (**Appendi**x **B.1**). Thus, there is a need for fully unsupervised methods capable of leveraging the underlying gene-level information to identify interaction-specific gene programs.

Several fully unsupervised methods exist for quantifying tissue regions. Implicit within each of these methods is the representation of regions in terms of low-frequency (i.e. large-scale) gene expression patterns over the tissue (Pham et al., 2023; Hu et al., 2021; Xu et al., 2024; Dong & Zhang, 2022; Long et al., 2023; Varrone et al., 2024; Chidester et al., 2023; Hu et al., 2024). These patterns can be isolated via low-pass filtering, i.e. by “smoothing” gene expression across neighboring cells (**Fig. 1a,b**, top) (Chang et al., 2022; Ortega et al., 2018; Wu et al., 2019; Zhang et al., 2015). The resulting patterns can then be grouped into region-specific gene programs via PCA (**Fig. 1c**, top), and cells can be clustered on the basis of these programs to identify regions. This approach is quantitatively appealing in that it leverages the underlying gene-level information to identify interpretable region-specific gene programs. However, it does not directly provide insight into interactions.

**Figure 1.**
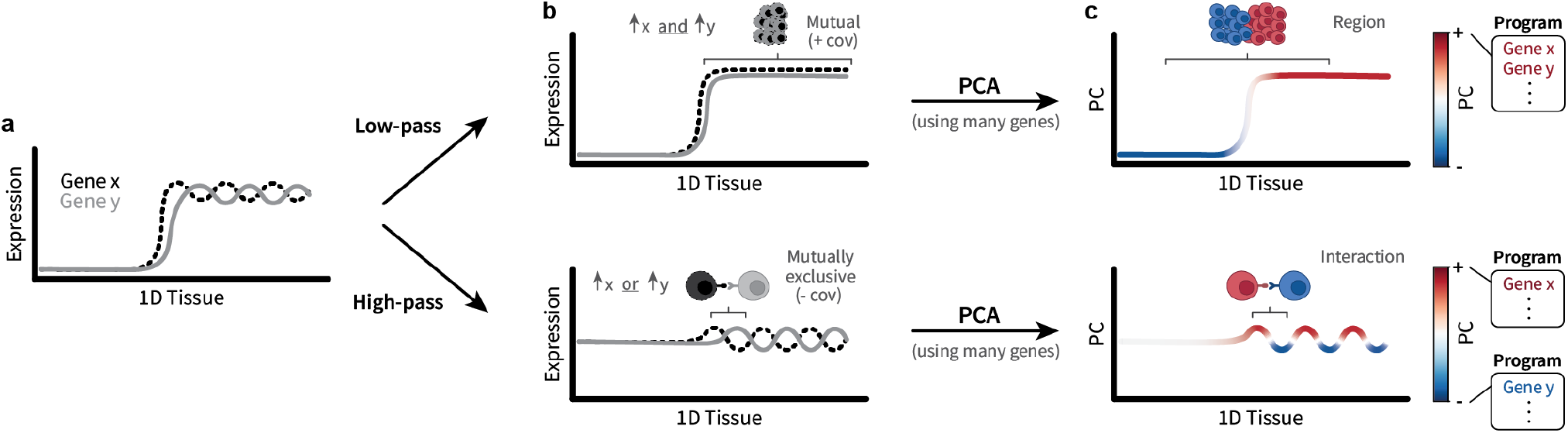
Illustration of the proposed framework. **a)** Two gene expression signals over a one-dimensional tissue. One can imagine moving left and right through the tissue, i.e. the x-axis. **b)** Positive/negative covariance (“cov”) between low-/high-frequency patterns represents regions/interactions. **c)** This covariance can be captured by PCA, allowing identification of gene programs underlying regions/interactions.

Here, we extend this frequency-based (i.e. harmonic) framework to interactions. Just as regions are represented by *positively* covarying *low*-frequency gene expression patterns over the tissue, interactions can be represented by *negatively* covarying *high* frequencies (**Fig. 1b**, bottom). The resulting covariance can then be decomposed into interaction-specific gene programs via PCA (**Fig. 1c**, bottom). Furthermore, we show that low and high frequencies can be merged to create mid frequencies, which can be interpreted as “interactions between regions” or region boundaries. This generalizes interactions to larger length scales, which can be equivalently interpreted as paracrine interactions. Finally, the resulting equations reveal a duality in which regions and interactions are two alternate representations of the same underlying information, each with unique strengths and weaknesses; regions are robust yet lack mechanistic insight, while interactions provide such insight at the cost of robustness.

We first demonstrate each concept in a 10X Xenium dataset from human lymph node (10X Genomics, 2024c). Low frequencies reveal canonical lymph node anatomy, high frequencies reveal T-cell activation by dendritic cells, and mid frequencies reveal T-cell migration patterns from blood vasculature into the surrounding tissue. We then recapitulate each result in an additional Xenium dataset from human tonsil – a similar secondary lymphoid tissue (10X Genomics, 2024a). Finally, to determine the extent to which this approach generalizes across experimental technologies, species, and tissue types, we identify disease-associated regions and interactions in a STARmap dataset from Alzheimer’s disease mouse model brains (Wang et al., 2018) and validate each result in an analogous Xenium dataset (10X Genomics, 2024b).

Supplementary figures, mathematical details, conventional cell typing results for each dataset, and connections to image processing and graph neural networks can be found in the **Appendi**x. Notebooks and scripts for reproducing the figures shown in this manuscript can be found at:

github.com/wanglab-broad/harmonics

## 1. Preliminaries

The proposed framework for defining and identifying regions and interactions relies on filtering. Filtering is the deliberate removal of certain signal components in order to emphasize others. For example, boosting the bass in a song corresponds to increasing the lower frequencies, thereby relatively decreasing the higher frequencies. In the time domain, over which such auditory signals occur, we are given a “frequency basis” composed of sinusoids of varying frequencies that, when combined in the right amounts, can represent any sound. However, spatial transcriptomics consists of gene expression signals over a relatively complex spatial *graph* domain that represents the tissue. In this graph domain, we are not simply given a frequency basis; we must calculate it. Here, we illustrate this calculation and how it enables filtering of gene expression signals over tissue domains.

### 1.1 Gene expression signals over tissue domains

Spatial transcriptomics data can be represented as a set of molecular signals over a spatial tissue domain. The tissue domain is represented by an undirected graph. Connectivity is calculated using a Delaunay triangulation and represented by the symmetric adjacency matrix **A** ∈ {0, 1}^*n*×*n*^. Each entry of **A** is either 1, which represents two cells that are connected, or 0, which represents two cells that are not connected. The number of connections for each cell *i* is given by the diagonal degree matrix **D** ∈ ℝ^*n*×*n*^ with entries **D**_*ii*_ = Σ_*j*_ **A**_*ij*_. A gene expression signal over the tissue is defined as a vector x ∈ ℝ^*n*^, with each element x_*i*_ representing the amount of expression in cell *i*. The full cell-by-gene matrix is then represented as a collection of signals, **X** = [x_1_|…|x_*g*_] ∈ ℝ^*n×g*^, where each column represents a gene signal, and *g* is the number of genes measured.

### 1.2 Frequencies

Given the above definitions, we can now calculate a frequency basis over the tissue domain which will be used to filter gene expression signals. Note that the underlying mathematics shown here corresponds to existing fundamentals of graph signal processing (Ortega et al., 2018).

The notion of frequency corresponds to a change in signal as one moves through a domain. Low-frequencies entail small changes as one moves along the domain, while high-frequencies entail large changes. Thus, given a signal and a domain, we can quantitatively describe this notion of frequency. One can picture moving along a graph domain as taking a step between nodes *i* and *j*. The change in a signal **v** during that step is given by **v**_*i*_ − **v**_*j*_. As we only care about the absolute change, we can square it to get (**v**_*i*_ − **v**_*j*_)^2^. To calculate an overall frequency value of **v** over the whole tissue, we can average all of the squared differences observed over the entire graph:

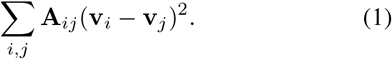

Note that while the sum is over all pairs of cells *i, j*, multiplying plying by the corresponding entries of the adjacency matrix **A**_*ij*_ ensures we only consider differences between neighboring cells. In the literature, eq. (1) is occasionally referred to as the “total variation” or “Dirichlet energy” of **v**.

We now have a metric that represents the frequency of a given signal. However, our goal is to construct a basis of multiple “pure” sinusoidal signals of varying frequency, analogous to those given by the time domain. To calculate this basis, first consider that eq. (1) can be expressed alternatively as the quadratic form **v**^⊤^**Lv**, where **L** = **D** − **A** is the graph Laplacian matrix (**Appendi**x **B.2**). Signals with minimal and maximal frequencies can then be represented as Rayleigh quotients:

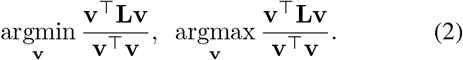

A set of unique solutions for expressions of this form is given by the eigenbasis of orthogonal eigenvectors of **L**. This can be shown via the same Lagrangian-or Rayleigh quotient-based approaches often used to derive PCA (Ghojogh et al., 2023). We denote this eigenbasis by **V** = [**v**_1_|…|**v**_*n*_] ∈ ℝ^*n*×*n*^, where **v**_1_ and **v**_*n*_ correspond to the minimal and maximal frequencies described in eq. (2) and **v**_*i*_ for 1 *< i < n* correspond to frequencies in between. We visualize some of these frequencies in **Fig. 2b**. The values associated with each frequency are given by the corresponding eigenvalues, *λ*_1_, …, *λ*_*n*_. Note that no measured gene expression information has been used up to this point. Rather, this basis represents a collection of “ideal” frequencies, analogous to sine waves in the time domain, which are inherent to the domain itself.

**Figure 2.**
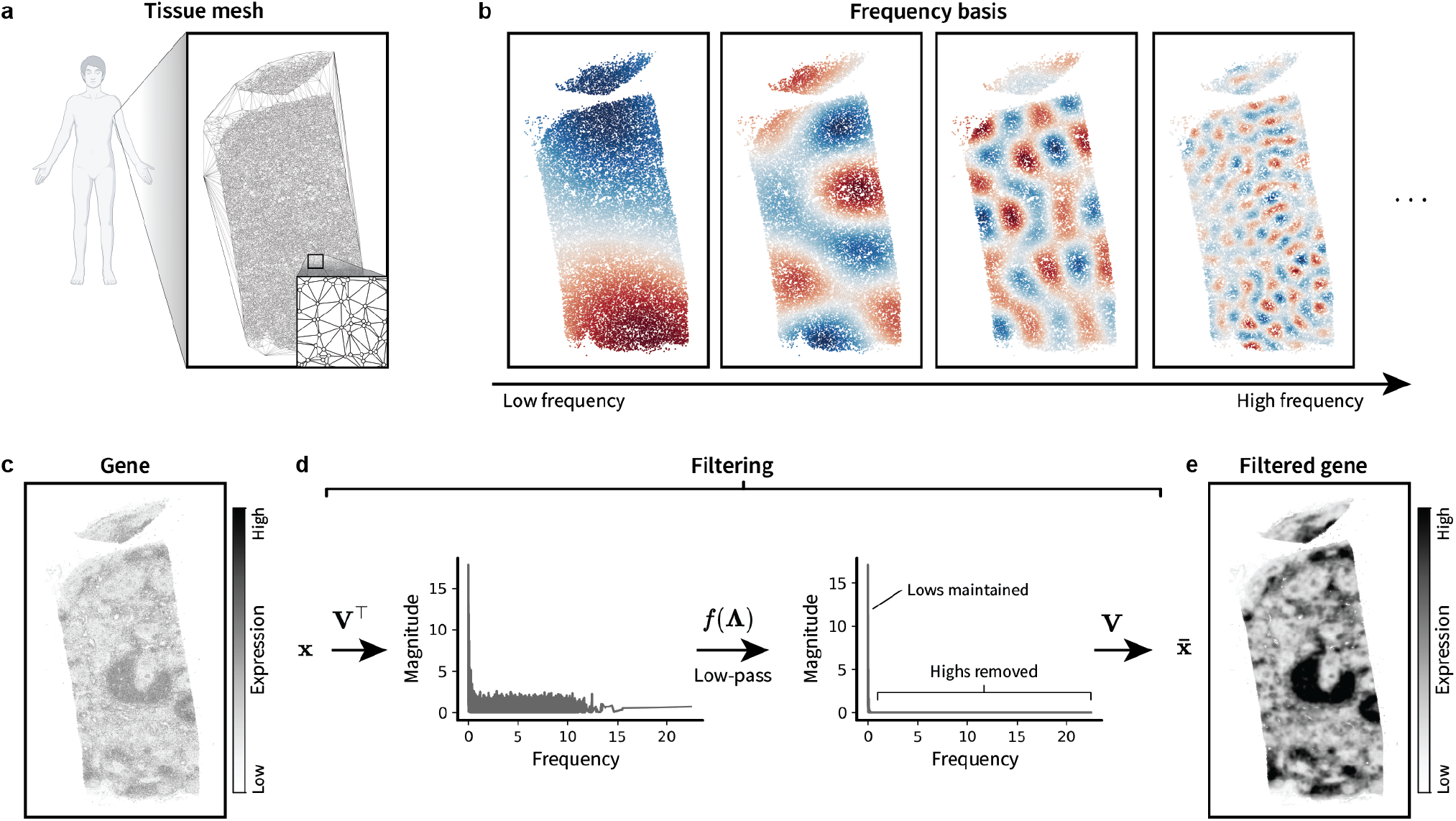
**Demonstration of filtering** in a 10X Xenium dataset from a human lymph node. Data contains 377 genes measured over 377,985 cells. **a)** The spatial tissue graph. Each node represents a cell. Edges are calculated using a Delaunay triangulation. **b)** Example frequencies plotted over the tissue. **c-e)** Visualization of low-pass filtering (i.e. eq. (3)). **d)** For each curve, each point represents a weight for a given frequency, such as those visualized in b). The gene pictured is *TRAC*.

At this point, one might ask “Why only *n* frequencies?” or “Is *n* enough?” The answer lies in the fact that the matrix **V** is an eigen*basis*, which means it is composed of *n* orthogonal unit vectors and is thus sufficient to represent any *n*-dimensional signal. Therefore, we can take any gene expression signal x ∈ ℝ^*n*^ (**Fig. 2c**) and project it onto this frequency basis via **V**^⊤^x (**Fig. 2d**, left), which corresponds to the Fourier transform. This can be interpreted as calculating how similar the signal x is to each frequency **v**_*i*_, yielding a “spectral” representation of x. Intuitively, this spectrum describes a recipe of how much of each frequency is required to construct x.

### 1.3 Filtering

Filtering entails modification of this spectral recipe. To filter the signal x, we modify the gene’s spectrum via point-wise multiplication with a kernel function, *f* . This kernel is thus a function over the Laplacian eigenvalues (i.e. frequency values). For example, a low-pass kernel, such as the diffusion kernel *f* (*λ*) = *e*^−*τλ*^, enhances low frequencies based on the parameter *τ*, with higher values of *τ* enhancing lower frequencies. Point-wise multiplication yields the modified spectrum *f* (**Λ**)**V**^⊤^x, where **Λ** is the diagonal matrix of Laplacian eigenvalues (**Fig. 2d**, right). This modified spectrum can then be transformed back into the tissue domain by multiplying by the inverse of the frequency basis (**Fig. 2e**). As the frequency basis is an orthogonal matrix, its inverse is simply its transpose. Thus, altogether, this filtering process corresponds to the equation

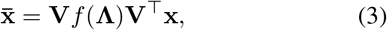

where 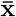 is the resulting low-pass filtered gene expression signal over the tissue. For the demonstration in Fig. 2, we used a diffusion kernel with *τ* = 5.

A related observation is that any function of the Laplacian matrix describes a filter. To see this, first diagonalize the Laplacian matrix as **L** = **VΛV**^⊤^. Any function of the Laplacian is a function of its eigenvalues, as its eigenvectors simply provide a change of basis:

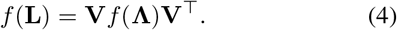

Comparing with eq. (3), one can see that 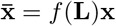 . Thus, the matrix *f* (**L**) corresponds to a filter with kernel *f* . We will use this property in Section 3 to express interactions in terms of high frequencies.

While the above mathematics provides the theory behind filtering, approximations are typically used in practice due to the high computational cost of eigendecomposition. In fact, the explicit eigendecomposition necessary to visualize the frequency basis and spectra shown in Fig. 2b,d had to be calculated using a random ∼ 10% subset of the data. However, all other results in this manuscript were calculated using efficient approximations implemented in the Python package `PyGSP` (Hammond et al., 2009; Defferrard et al., 2017), enabling efficient analysis of the full dataset (377 genes measured across 377,985 cells).

With the above understanding of filtering, we can now derive representations of regions and interactions.

## 2. Regions

### 2.1 Representation

As mentioned above, recent literature implicitly represents regions in terms of groups of positively covarying low-frequency gene expression signals (**Fig. 1**, top). In other words, two low-pass filtered (and mean centered) genes, 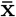 and 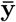, represent a region if

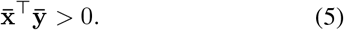

We can generalize this equation to all pairs of genes by substituting 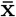 and 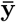 each with the full cell-by-gene matrix of filtered and mean centered gene signals,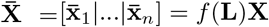 . This yields the low-pass covariance matrix 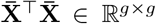, which forms a table where each entry contains a comparison of two genes in the form of eq. (5). Eigendecomposition of this covariance matrix simply corresponds to PCA, in which genes are grouped into programs based on their large-scale coexpression patterns. Cells are then represented in terms of these programs in PC space (**Fig. 1c**, above). Finally, clustering can be performed in PC space to calculate region labels for each cell. Note that the above process simply corresponds to conventional single-cell analysis with the addition of an initial low-pass filtering step.

### 2.2 Demonstration

We demonstrated region identification in the same lymph node dataset introduced in Section 1. Using `Scanpy` (Wolf et al., 2018), we preprocessed the raw read counts data by normalizing to 10,000 counts per cell (i.e. `sc.pp.normalize total`) and transforming with a shifted logarithm (i.e. `sc.pp.log1p`). We then filtered using the diffusion kernel mentioned above, scaled the filtered signals to unit variance and zero mean (i.e. `sc.pp.scale`), and clipped values to *±*2 standard deviations (**Fig. 3a**). We then performed PCA and k-means clustering using `scikit-learn` (Pedregosa et al., 2011) (**Fig. 3b,c**). We chose k-means for its simplicity as well as to avoid smoothing-induced artifacts found in nearest-neighbors-based methods (e.g. Leiden) (Maher et al., 2023). We chose *k* = 3 clusters to reflect the main region features observed in top PCs as well as basic lymph node anatomy (Willard-Mack, 2006).

**Figure 3.**
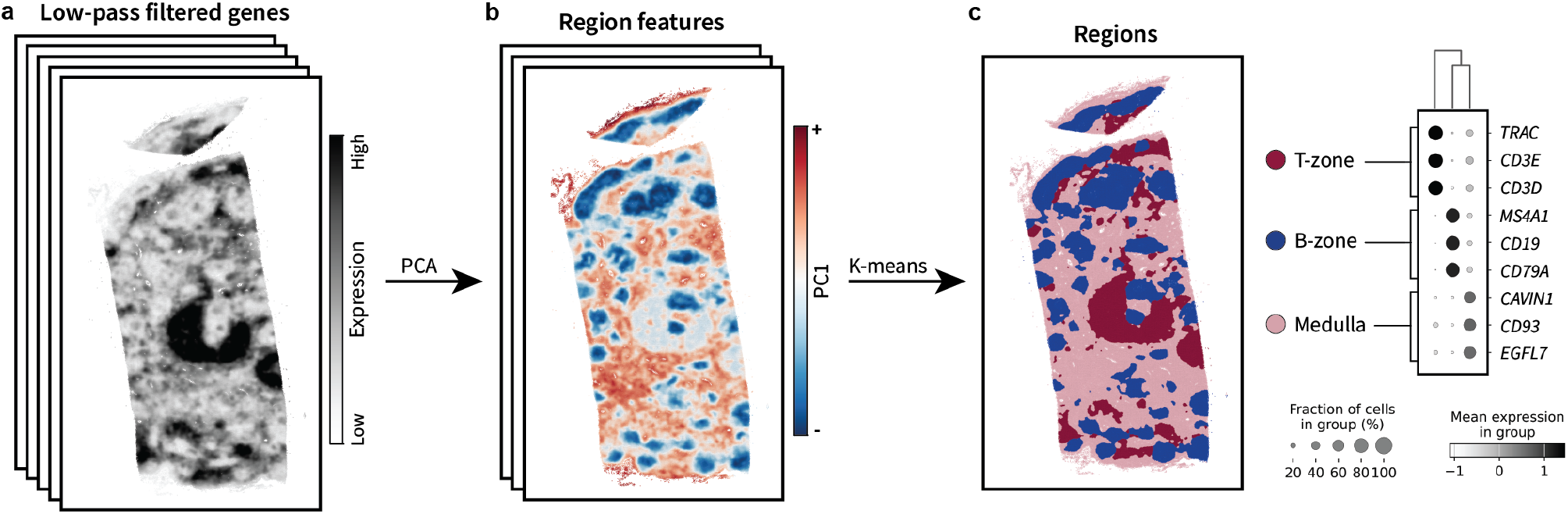
**Demonstration of region identification** in the human lymph node. **a)** Low-pass filtered gene expression signals (i.e. output shown in Fig. 2e). **b)** The resulting PC1. **c)** Region clusters calculated using k-means alongside their corresponding marker genes.

The resulting regions indeed reflected the conventional anatomy, with B-cell zones (“B-zone”) resembling the cortex, T-cell zones (“T-zone”) resembling the paracortex, and the medulla defined by connective and vascular tissue markers. Yet, just as in the existing region identification methods mentioned above, further analysis would likely be required to link these regions to more mechanistically informative features, such as interactions.

## 3. Interactions

### 3.1 Representation

Cell-cell interactions often display coordinated transcriptional patterns between neighboring cells. For instance, mutually exclusive expression patterns may be required to produce complementary surface proteins that enable binding between two cells (Schenkel & Pauken, 2023). On the other hand, an existing interaction may lead to downstream transcriptional changes that are unique to each cell. In either case, each interacting cell would likely display a unique transcriptional signature when engaged in an interaction. It is this fundamental assumption of mutually exclusive gene expression patterns between interacting cells that we use to derive a quantitative representation of interactions.

We can first gain an intuition for this pattern by visualizing an instance of mutually exclusive gene expression in two neighboring cells, *i* and *j*:

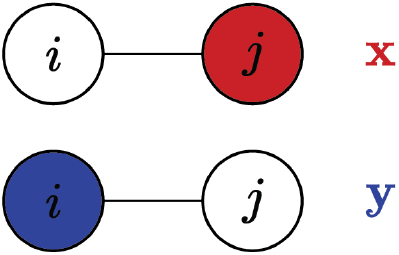

For an interaction to occur between these two cells, one cell must express gene *x* (red) but not gene *y* (blue), while the other cell must express *y* but not *x*. Additionally, we expect to observe this pattern frequently within the tissue. We can translate these pieces of intuition into a simple equation. First, consider the differences in gene expression between each cell, (x_*i*_ − x_*j*_) and (y_*i*_ − y_*j*_). Based on the above diagram, we would expect the first difference to be a large positive value and the second a large negative value. Thus, if we multiplied these differences, we would overall expect a large negative value: (x_*i*_ − x_*j*_)(y_*i*_ − y_***j***_) *<* 0. Finally, given that we expect to observe this interaction pattern often within the tissue, we could calculate this product for all pairs of cells and take the sum, weighting by the adjacency matrix to consider only neighboring cells. Altogether, this describes the following equation:

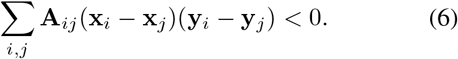

This equation can alternatively be expressed in terms of frequencies. Note that it appears to be a generalization of eq. (1) to two different genes. Indeed, just as eq. (1) can be expressed as the quadratic form x^⊤^**L**x, so can eq. (6) be expressed as x^⊤^**L**y (**Appendi**x **B.3**). The Laplacian can then be split into two terms, yielding 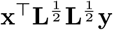 and further 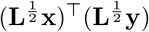 . By eq. (4) (i.e. “any function of the Laplacian is a filter”), we can see that 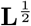 represents a square root filter with kernel 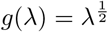, which is high-pass due to the square root function’s higher weighting of higher values. We can then express our high-pass filtered gene expression signals as 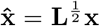 and 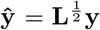, ultimately leading us to a representation of interactions as

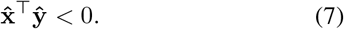

Assuming mean-subtraction after filtering, this equation states that interactions are represented by negatively covarying high-frequency components of genes *x* and *y*.

Just as for low-pass filtering, we can independently high-pass filter each gene to obtain the filtered cell-by-gene maTrix 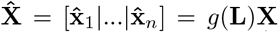, where 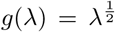. The high-pass covariance matrix 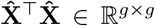 can then be eigendecomposed to identify gene programs underlying interactions (i.e. PCA on 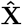) (**Fig. 1c**, below). However, unlike in the low-pass case, we expect interacting gene programs to separate onto *opposite* sides of a given PC because they *negatively* covary. As a result, we explicitly do not want to cluster. Rather, we interpret each PC as a unique interaction between cells expressing either the positive or negative gene program.

### 3.2 Demonstration

We next sought to demonstrate this interaction identification approach using the same human lymph node dataset introduced above. One of the most prominent interactions that takes place within in the lymph node is T-cell activation, which occurs between T-cells and dendritic cells in the T-zone. Thus, we attempted to leverage the region clustering results from Section 2 to identify T-zone-specific interactions.

We first computed the high-pass covariance between all cells within the T-zone only, denoted 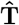 (**Fig. 4a,b**). Next, we separately calculated the covariance among all cells *outside* the T-zone, denoted by **Ŝ**, and divided it out from 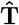. Conceptually, this corresponds to finding transcriptional relationships between adjacent cells in the T-zone (i.e. 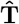) and then removing (i.e. dividing out) relationships found outside the T-zone (i.e. **Ŝ**). Scalar division corresponds to multiplying by an inverse, and this holds for matrices as well. Thus, this new T-zone-specific covariance matrix can be expressed as 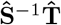. Just as before, 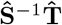 can be eigen-decomposed to identify interaction-specific gene programs. This eigendecomposition simplifies to 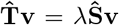, yielding a generalized eigenvalue problem for which standard solvers exist (**Appendi**x **B.4**) (Virtanen et al., 2020). Note that by setting **Ŝ** = **I**, we get a standard eigenvalue problem (corresponding to standard PCA on 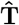).

**Figure 4.**
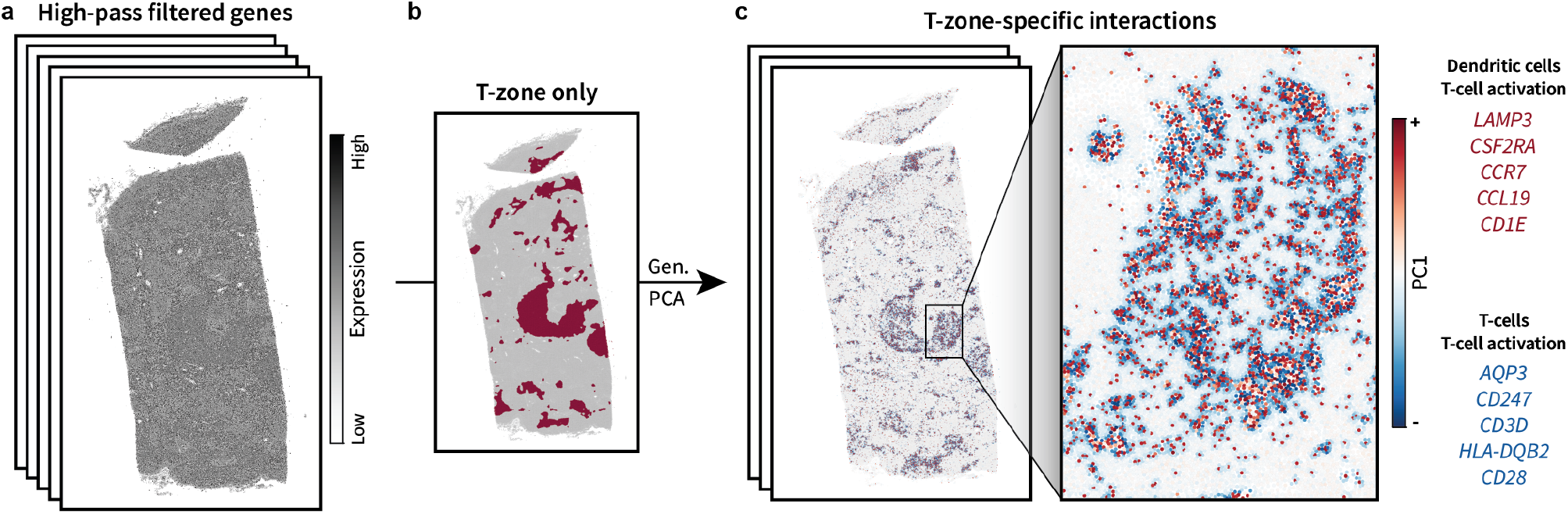
**Demonstration of region-specific interaction identification** in the human lymph node. **a)** High-pass filtered gene expression signals. **b)** Isolation of the T-zone (dark red) in order to consider interactions only within this region. A “generalized” (Gen.) PCA is performed by additionally dividing out patterns observed in the rest of the tissue (light grey). **c)** The resulting PC1 plotted in the tissue. The top 5 markers for each side of PC1 (i.e. each interaction program) are listed in descending order of relative contribution.

Upon using this T-zone-specific “generalized PCA” approach, the resulting PC1 revealed strongly localized patterns between two cell populations within the T-zone (**Fig. 4c**). The corresponding interaction-specific gene programs indicated that the cell population on the positive side of PC1 represented dendritic cells, while cells on the negative side were T-cells. This was reflected in conventional cell typing results (**Fig. A6**). However, perhaps most importantly, the relative contribution of each individual gene within each program enabled interpretation of cell *states*. For instance, note that the top gene in the positive program was *LAMP3*. This is a known marker of mature dendritic cells enriched in immunoregulatory molecules (mregDCs), which have a particularly high propensity for engaging in immunoregulatory interactions (Li et al., 2023). Thus, negatively covarying high frequencies indeed appeared to represent interactions.

We found that the generalized PCA approach was necessary to identify interactions in this dataset. Calculating standard PCA using 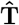 (i.e. not dividing out **Ŝ**) yielded noisy results (**Fig. A1**). In particular, positive and negative gene programs did not appear to be highly expressed between adjacent cells. Rather they were sparsely scattered across the entire tissue and described generic cell types. The same was true when calculating interactions across the entire tissue (i.e. not restricting our search to the T-zone) (**Fig. A2**). We hypothesized that the reason for this poor performance was that eq. (6) is essentially an average over the entire tissue. As a result, interactions that occur only within a subset of the tissue might be averaged away. However, the generalized eigendecomposition approach strongly focuses analysis on a particular region while distinguishing it from all others, thus providing sufficient specificity so as to avoid this “averaging out” problem.

## 4. Boundaries

### 4.1 Representation

Having interpreted low frequencies as regions and high frequencies as interactions, we can now turn to the frequencies in between. Such “mids” can be expressed as combinations of lows and highs, creating filters that only maintain the shared frequencies toward the middle. Conceptually, one can think of this as applying a preliminary low-pass filter to find region features followed by a high-pass filter to find interactions between them. Thus, mids can be interpreted as “interactions between regions” or, equivalently, region boundaries. More generally, however, they can be thought of as an extension of interactions to larger length scales.

We can make this intuition quantitatively explicit. Consider two gene expression signals, x and **y**. These signals can be sequentially low-pass and then high-pass filtered using the previous low-pass kernel from Section 2.1, *f* (*λ*) = *e*^−*τλ*^, followed by the high-pass kernel from Section 3.1, 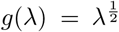. This yields the “mid-pass” filtered signals *g*(**L**)*f* (**L**)x and *g*(**L**)*f* (**L**)**y**. As in previous sections, we consider their relationship by computing their covariance, x^⊤^*f* (**L**)*g*(**L**)*g*(**L**)*f* (**L**)**y**. Setting 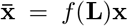 and 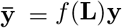, this simplifies to

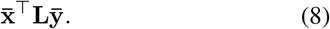

Note that this is the same as eq. (7) but with 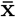 and 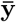 substituted for x and **y**. Thus, it indeed represents interactions between region-level features.

Alternatively, the above equations can be rearranged and interpreted in terms of paracrine interactions. The previous section described juxtacrine interactions, in which cells communicate via direct contact of molecules attached to their surfaces (Perrimon et al., 2012). Paracrine interactions, on the other hand, are defined as communication between cells along larger length scales via *diffusion* of secreted signaling molecules from one cell to another. We can express this quantitatively by first noting that *f* (**L**) and *g*(**L**) are commutative, as they are both functions of the Laplacian matrix and thus share the same eigenbasis, as seen via eq. (4). In terms of linear algebra, this corresponds to simultaneous diagonalizability. In terms of signal processing, this corresponds to the convolution theorem. Because these matrices are commutative, we can rearrange them to move all low-pass filtering matrices to the left, yielding x^⊤^*f* (**L**)*f* (**L**)*g*(**L**)*g*(**L**)**y**. Setting 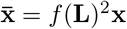, this simplifies to

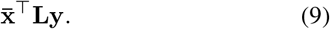

This can be interpreted as applying a (more aggressive) lowpass filter to signal x – effectively modeling the diffusion of corresponding signaling molecules from “sender” cells – and then calculating the amount that it interacts with **y** in “receiver” cells. Thus, negatively covarying mids can also be interpreted as paracrine interactions.

Note that eqs. (8) and (9) can also be expressed as summations in the form of eq. (6). For instance, one can expand eq. (9) to obtain

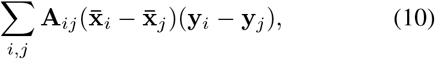

providing a bridge to the neighborhood-level intuition given in the previous section.

### 4.2 Demonstration

To demonstrate this representation in the human lymph node, we first performed mid-pass filtering (**Fig. 5a**). We then performed PCA to identify boundary-specific gene programs. The negative gene program associated with PC1 appeared to represent blood vasculature, as it was increased within circular openings in the tissue that resembled vessels and consisted of markers such as *PECAM1* and *CD34* (**Fig. 5b**). The positive gene program formed a boundary with the vasculature, containing T-cell markers such as *TRAC* and genes involved in T-cell chemotaxis such as *PTPRC* and *CXCR4* (Fernandis et al., 2003). Furthermore, the morphology of the observed vasculature appeared similar to that of high endothelial venules (HEVs), and interactions between the products of *CD34* and *SELL* are known to mediate T-cell migration out of HEVs and into the lymph node (Blanchard & Girard, 2021). Thus, while this could be viewed as a *boundary* between vasculature and neighboring areas enriched in migrating T-cells, it could also be interpreted as a chemotactic *interaction* occurring over a larger length scale. Finally, note that additional PCs appeared to show boundaries between the interior and exterior of germinal centers (**Fig. A3**). Altogether, it appeared that negatively covarying mids represented interactions on larger length scales.

**Figure 5.**
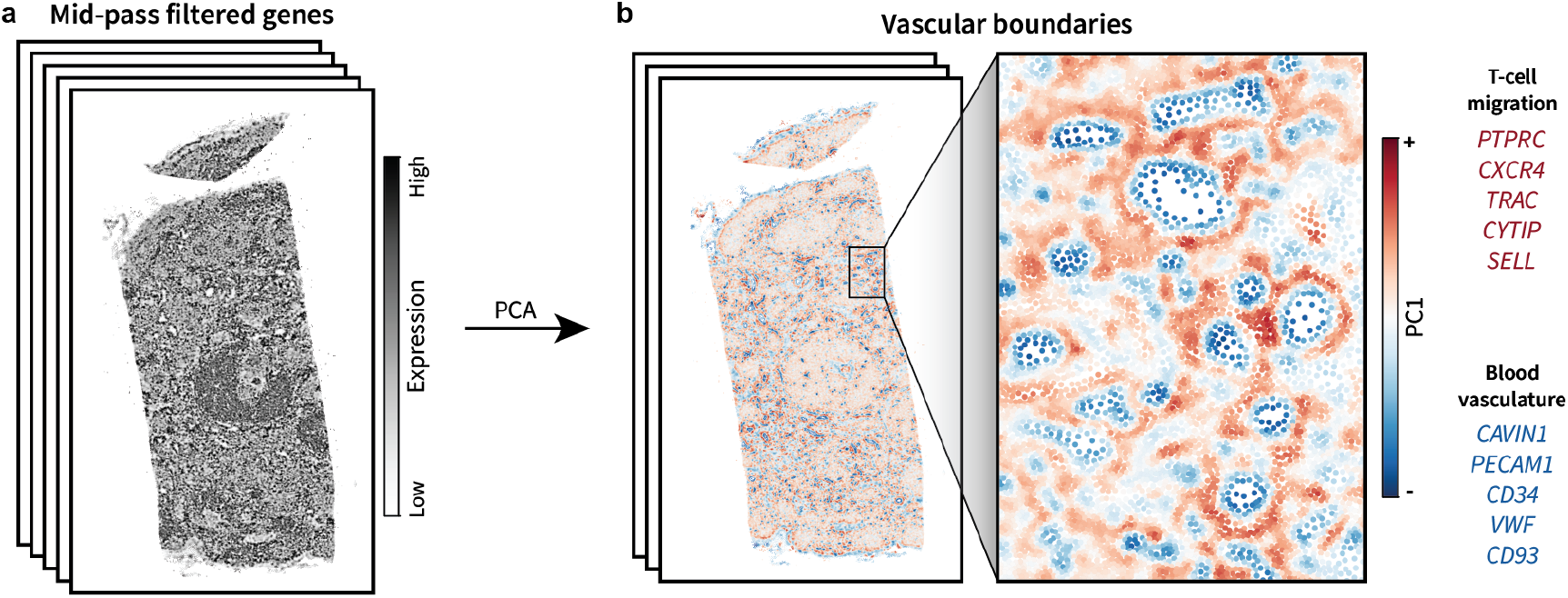
**Demonstration of boundary identification** in the human lymph node. **a)** Mid-pass filtered gene expression signals. **b)** The resulting PC1 plotted in the tissue. The top 5 markers for each side of PC1 are listed in descending order of relative contribution.

## 5. Duality

It may seem thus far that interactions provide more mechanistic insight than regions. Indeed, the positive and negative sides of interaction-specific programs might be interpreted as the attractive molecular “forces” shaping tissue structure and function, while regions might be interpreted as the result of such forces. However, regions and interactions are nearly equivalent concepts. On the one hand, a region could be defined as an area in which certain interactions are likely to occur. On the other hand, interactions require spatial proximity between cells, constituting aggregates that could instead be viewed as regions. In this section, we show that this conceptual “duality” can be expressed quantitatively. This allows us to explicitly derive unique strengths and weaknesses of each representation, ultimately suggesting that interactions indeed convey more information than regions yet are less robust.

### 5.1 Representation

Representations of regions and interactions are determined by their corresponding filters. For instance, consider the high-pass filter used to define interactions above:

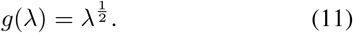

To create an equivalent region representation, we simply reverse this filter such that its preference for high frequencies becomes a preference for low frequencies (**Fig. 6b**). The reversed form of the above filter is

**Figure 6.**
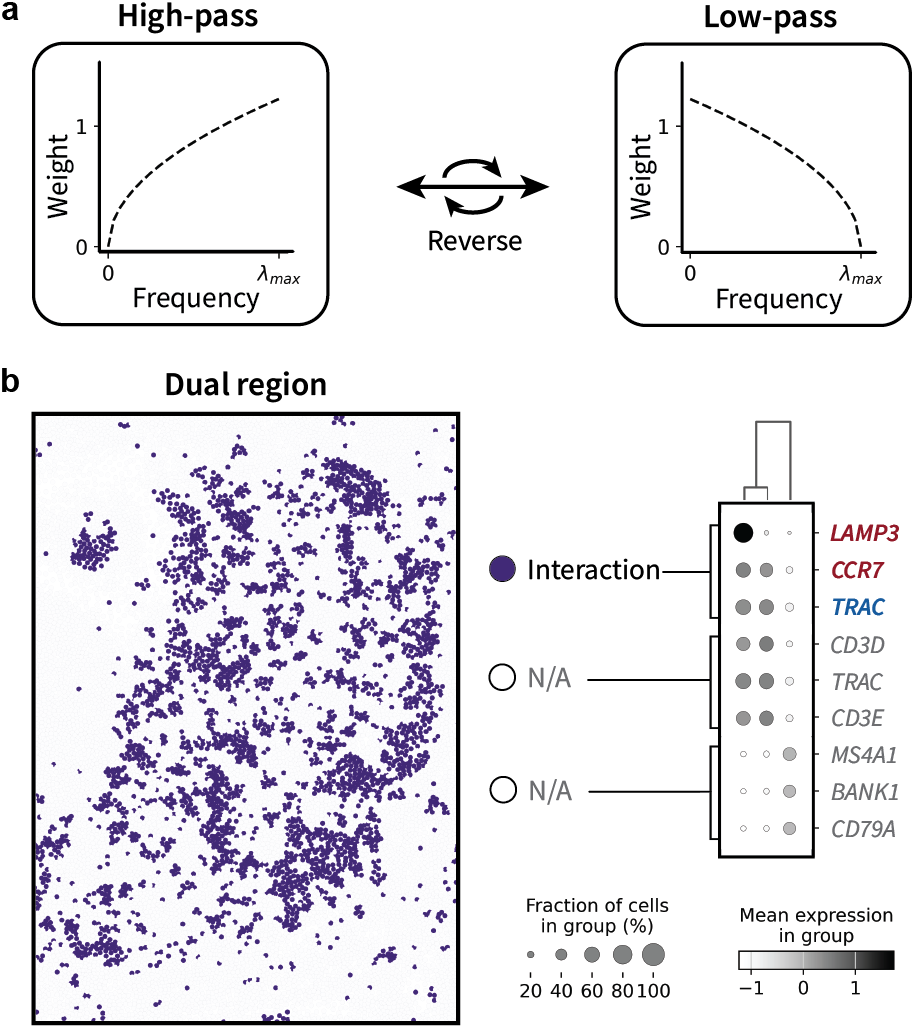
**Demonstration of duality between regions and interactions** in the human lymph node. **a)** Schematic of filter reversal. **b)** The dual region obtained from applying a reversed interaction filter followed by k-means. Gene markers for the purple region are colored red and blue to match their role in the interaction-specific gene program shown in Fig. 4c. Unrelated regions labeled “N/A”.

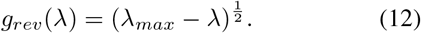

Intuitively, reversal should produce an “equally aggressive” low-pass filter such that the resulting regions exist on the same length scale as the original interactions. We can explicitly show that these length scales match by expressing each filter in terms of matrices and then summations.

To express reversal in terms of matrices, we first normalize the adjacency matrix for convenience: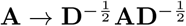.

The normalized Laplacian is then given by **L** = **I** − **A** with sorted eigenvalues *λ*_*min*_, …, *λ*_*max*_ ∈ [0, 2]. Next, recall from eq. (4) that any filter kernel *g*(*λ*) can also be understood as a matrix function, *g*(**L**). Applied to eq. (11), this yields

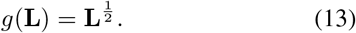

Similarly, the matrix function form of the reversed filter in eq. (12) is given by

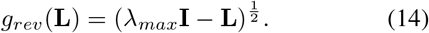

The matrix given by **Γ** = *λ*_*max*_**I** − **L** can be understood as somewhat of a “reversed Laplacian” in that it has the same eigenvectors as **L** but with reversed assignment of eigenvalues (**Appendi**x **B.5**). As a result, the reversed form of filter *g*(**L**) is simply *g*(**Γ**). Note that **Γ** is also similar to the “renormalized” and “signless” Laplacians (Wu et al., 2019; Kipf & Welling, 2017; Luan et al., 2022).

We can now use these filters to define regions and interactions in the style of eqs. (5) and (7). Just as in Section 3.1, we can define interactions by first performing high-pass filtering to obtain 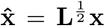 and 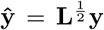 . Covariance between high frequencies is then given by

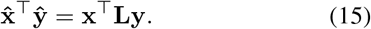

Similarly, for regions, we now have 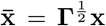 and 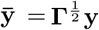. Covariance between low frequencies is then given by

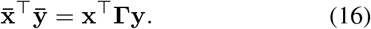

Finally, we can expand eq. (16) into a summation for interpretation. In Section 3.1, we began with the summation given by eq. (6) and simplified it to eq. (15). Here, we instead begin with eq. (16) and reverse this process (**Appendi**x **B.6**). If we assume, for simplicity, that *λ*_*max*_ = 2, we obtain

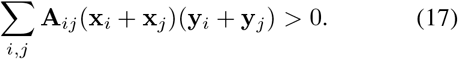

Note the similarity between eqs. (6) and (17): while the former represents interactions as neighborhood *differences*, the latter represents regions as neighborhood *sums*. This provides a quantitative rationale for the intuition that filter reversal should produce regions on the same neighborhood scale as their equivalent interactions.

### 5.2 Demonstration

We demonstrated this duality using the same human lymph node sample used in prior sections. We performed the same T-zone specific interaction identification as in Section 3.2 but with the reversed filter given by eq. (12) (**Fig. 6a**). We then performed k-means to cluster cells into regions, simply choosing *k* = 3 for consistency with our approach in Section 2.2. One of these clusters closely resembled the spatial pattern of the previously identified interactions (**Fig. 6b**). Furthermore, gene markers for this cluster overlapped with both the positive and negative interaction-specific gene programs, suggesting that it formed an aggregate representation of these interactions. For instance, *LAMP3* was a particularly strong marker of this cluster *alongside* T-cell-specific markers such as *TRAC*. Thus, it appeared that this region was indeed an aggregate of both the positive and negative interaction-specific gene programs found in Section 3.2.

### 5.3 Implications

Eqs. (6) and (17) reveal unique strengths and weaknesses of each representation. For instance, assuming no mean-centering, eq. (17) can only ever be positive, as all values within the summation are positive. This implies that regions offer a robust representation. However, they lack the ability to resolve different sides of an interaction because they simply aggregate features, decreasing their mechanistic interpretability. On the other hand, the differences in eq. (6) enable discrimination between different sides of an interaction. As a result, interactions offer more mechanistic interpretability than regions. However, they can result in both positive and negative values which “cancel out” during summation, leading to a less robust representation. Note that this is the same as the “averaging out” problem introduced in Section 3.2. Thus, each representation possesses unique strengths and weaknesses, with regions offering robustness and interactions offering interpretability.

These representations can also be interpreted as the two minimal modifications to the spatial covariance matrix such that it can be factorized into gene programs by PCA. As discussed in Appendix B.1, spatial covariance between gene signals x and **y** is typically defined as x^⊤^**Ay** (Chen, 2015; Miller et al., 2021; Palla et al., 2022b; Russell et al., 2023). However, this matrix is indefinite and thus has a mixture of positive and negative eigenvalues. As a result, it cannot be factorized by PCA. Now consider eqs. (6) and (16), which can also be expressed as x^⊤^(**I** − **A**)**y** and x^⊤^(**I** + **A**)**y**. These equations are nearly the same as spatial covariance. However, in each case, addition of the identity matrix shifts the eigenvalues up such that they become non-negative, thus yielding a positive semidefinite matrix that can be factorized into gene programs. This can be explained more conceptually as follows. The equation x^⊤^**Ay** only compares gene expression across *different* cells – it does not consider coexpression within the *same* cell. Yet gene programs are defined as groups of genes that are often coexpressed within the same cell. The addition of **I** simply allows one to calculate such *intra*cellular coexpression and thus obtain gene programs, while the addition or subtraction of **A** simultaneously incorporates *inter*cellular coexpression between neighbors. Thus, combining **I** and **A** allows representation of gene programs (intra) underlying colocalization between cells (inter). Whether **A** is added or subtracted determines whether these neighbors are aggregated or discriminated between.

## 6. Validation: human tonsil

In an effort to validate the above results, we repeated each analysis in a 10X Xenium dataset from a human tonsil with reactive follicular hyperplasia (10X Genomics, 2024a). As lymph nodes and tonsils are both secondary lymphoid tissues, we expected them to contain largely the same regions and interactions (van de Pavert & Mebius, 2010). Additionally, the tonsil dataset consisted of the same panel of 377 genes from the lymph node dataset, measured over 1,349,474 cells.

We first repeated region identification. We identified the same canonical secondary lymphoid anatomy comprised of T-zones, B-zones, and the medulla, all defined by similar marker genes (**Fig. 7a**). We also detected an additional region surrounding each B-zone that contained markers of cellular proliferation such as *SOX2* (Mamun et al., 2020). This likely represented the follicular hyperplasia corresponding to the sample condition. Aside from this region, however, the same canonical anatomy was indeed identified. Next, we repeated the T-zone-specific interaction analysis. Just as in the lymph node, PC1 identified colocalization between a positive gene program consisting of dendritic cell markers and a negative program consisting of T-cell markers (**Fig. 7b**). Furthermore, it showed *LAMP3* as the top marker for the positive program, again suggesting that these cells were participating in T-cell activation. We then repeated boundary identification and found that PC1 recapitulated the previous relationship between migrating T-cells and blood vasculature, resulting in similar morphological features and gene markers (**Fig. 7c**). Note that further PCs also showed boundaries between hyperplastic regions and T-zones (**Fig. A4**). Finally, we again reversed the T-zone-specific interaction filter and performed clustering to identify a corresponding “dual” region (**Fig. 7d**). Taken together, it appeared these results were replicable across different datasets and tissues.

**Figure 7.**
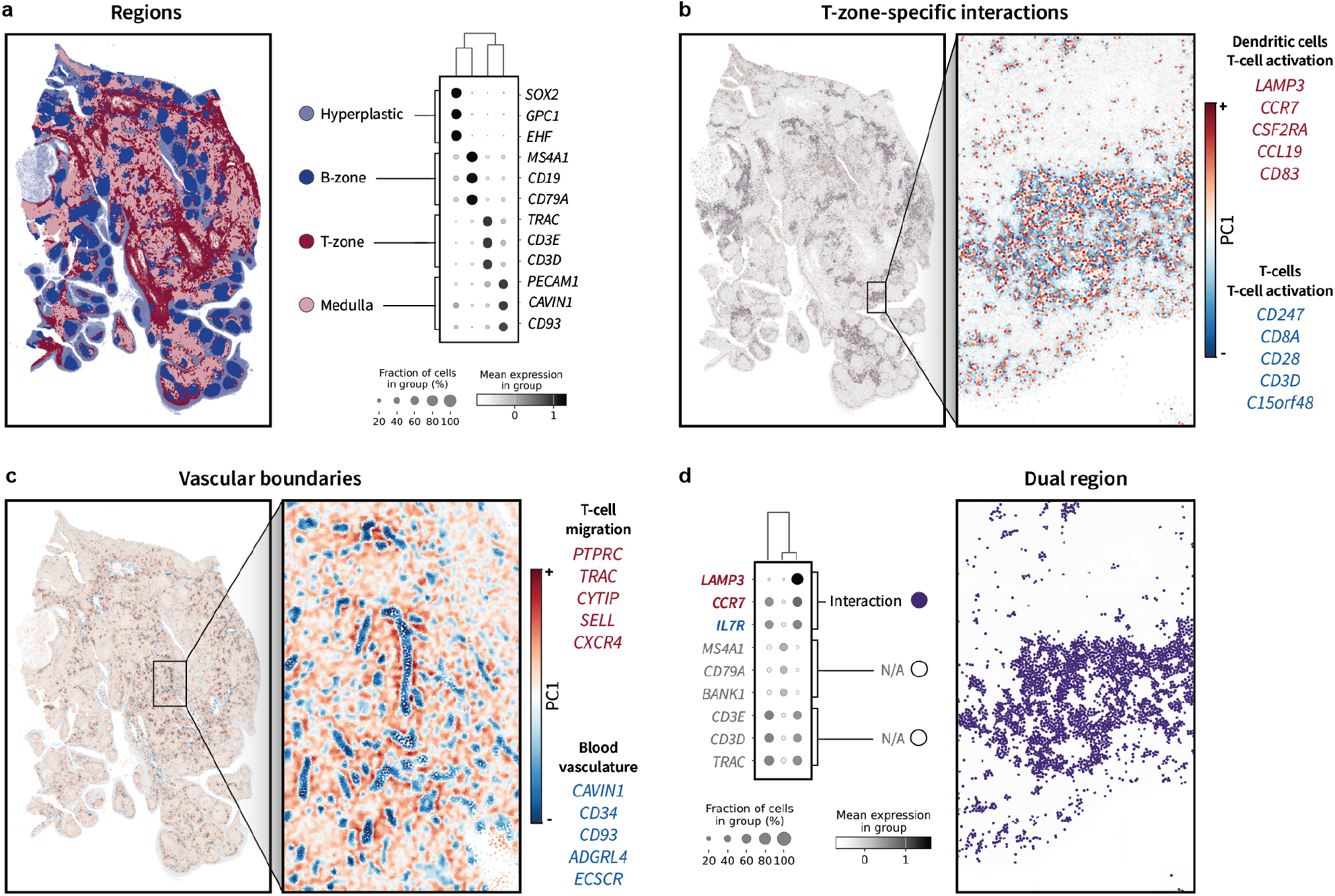
Recapitulation of above results in a 10X Xenium dataset from a human tonsil. Data contains 377 genes measured over 1,349,474 cells. **a)** Same as Fig. 3c. **b)** Same as Fig. 4c. **c)** Same as Fig. 5b. **d)** Same as Fig. 6b.

## 7. Application: mouse brain

We next sought to determine whether this framework would generalize to different experimental technologies and biological contexts. To do so, we turned to an existing Alzheimer’s mouse model brain dataset collected using STARmap (Zeng et al., 2023).

Alzheimer’s disease (AD) is a neurodegenerative disease characterized in part by the presence of extracellular amyloid-*β* plaques located throughout the hippocampus and neocortex. In this study, paired spatial transcriptomics and amyloid-*β* imaging was applied to control and disease mice at various ages to identify plaque-associated gene expression patterns. The data was analyzed by calculating cell type enrichment within various distance bins from each plaque. The authors found that disease-associated microglia (DAMs) were often located adjacent to plaques and surrounded by homeostatic microglia (HMs), suggesting that these microglial populations may interact with one another to influence disease progression. This hypothesis has since been supported by similar studies (Mallach et al., 2024).

However, the analyses in these studies required cell typing as well as binning into discrete distances from each plaque. We reasoned that harmonic representations might instead enable fully *unsupervised* detection of such interactions on the level of *genes*.

### 7.1 Demonstration: STARmap

The corresponding dataset contained multiple samples and multiple conditions. We performed filtering on each sample separately, as spatial coordinates in different samples were independent. We reasoned that, given similar spatial distributions of cells, frequency values should be comparable across different samples, and filtering should yield comparable results. After filtering, gene expression matrices for each sample were concatenated and analyzed together using PCA. Note that this was equivalent to conventional single-cell analysis of multiple samples with the exception of a preliminary filtering step. Finally, we accounted for control and disease conditions by leveraging the same generalized eigendecomposition method from Section 3.2, though to identify *disease*-rather than region-specific gene programs.

We first aimed to identify disease-specific region features using the dual region kernel given by eq. (12). The positive portion of the resulting PC1 formed small regions scattered across the brain in the same pattern as plaques shown in the original study (**Fig. 8a**). This pattern was restricted to the disease condition and increased with age. The gene program associated with this feature indeed consisted of DAM markers such as *Cst7, Clec7a*, and *Trem2* (Keren-Shaul et al., 2017; Deczkowska et al., 2018). The locations of these features also matched those of microglia obtained via conventional cell typing (**Fig. A8**). We then leveraged the paired amyloid-*β* stain by calculating distances between each cell in the disease condition and the nearest plaque, which decreased as PC1 value increased (**Fig. A5a**). Altogether, it appeared that low-pass filtering enabled unsupervised identification of plaque-associated regions.

**Figure 8.**
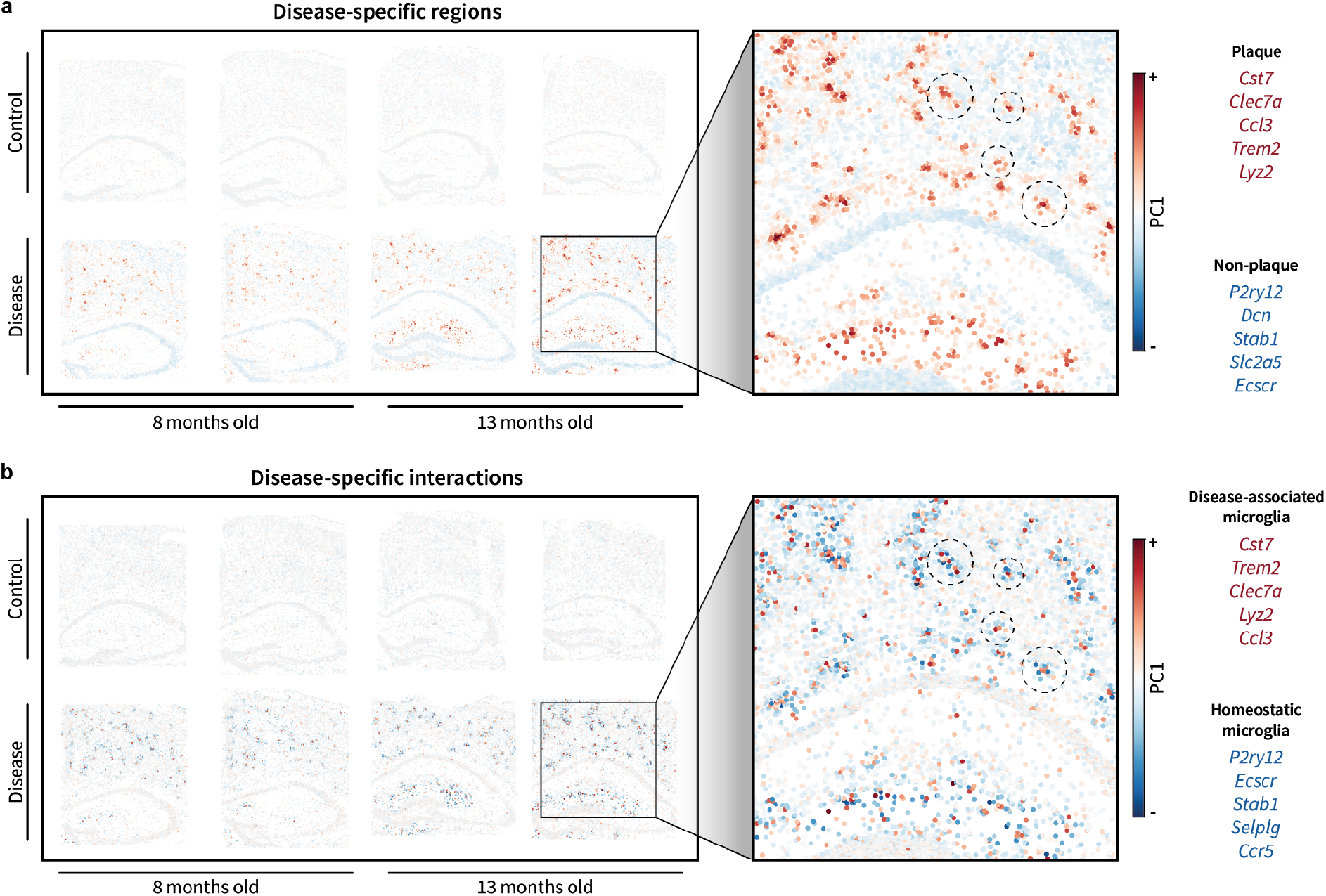
**Unsupervised identification of plaque-associated regions and interactions** in STARmap Alzheimer’s mouse model dataset. Data contains 2766 genes measured across 144,330 cells total. **a)** Results from applying the dual region filter to all samples followed by generalized PCA to identify disease-specific region features. The top 5 markers for each side of PC1 are listed in descending order of relative contribution. **b)** Same as a) but using the dual interaction filter.

However, the unique conceptual advantage of this approach is its ability to *decompose* such regions into potentially interacting cell populations, such as DAMs and HMs. To that end, we next filtered using the dual interaction kernel given by eq. (11). In absolute value, the resulting PC1 displayed the same overall spatial, disease-, and age-associated patterns as that of the dual region filter (**Fig. 8b**; note cells inside dotted circles). However, it showed additional variation in that the positive gene program (red) was expressed by cells in the center of each plaque region, surrounded by a shell of cells expressing the negative gene program (blue). Indeed, the positive gene program consisted of similar DAM markers, while the negative gene program consisted of HM markers such as *P2ry12* and *Selplg*. This pattern was also supported by the amyloid-*β* stain, as cells with positive PC1 values tended to be closer to plaques than cells with negative PC1 values (**Fig. A5b**). Thus, it appeared that this approach enabled fully unsupervised, gene-level characterization of the same putative DAM-HM interactions reported in the original study. Additionally, note that it identified the ligand-receptor pair *Ccl3* and *Ccr5*, which is known to play a role in immune signaling in the brain (Festa et al., 2023; Martin-Blondel et al., 2016).

### 7.2 Demonstration: Xenium

We again sought to validate our results in an additional dataset. We found that an analogous Alzheimer’s mouse model dataset was also available as a 10X Xenium demo (10X Genomics, 2024b). However, there were notable differences between these two datasets. They used different technologies, different mouse models, and different gene panels. Nonetheless, both technologies were single-cell resolved, both mouse models were plaque-forming, and both gene panels included microglial and signaling markers. Thus, we applied the same analysis approach as above and expected to observe similar results. Indeed, the PC1 resulting from low-pass filtering displayed a similar disease– and age-associated gene program corresponding to plaque regions (**Fig. 9a**). Additionally, the PC1 resulting from the dual high-pass filter appeared to similarly split each plaque region into a DAM-HM interaction program (**Fig. 9b**; note cells inside dotted circles). While the exact gene programs differed from the STARmap results, the biological meaning of each was largely the same. For instance, the positive program contained DAM markers such as *Itgax* and *Trem2*, while the negative program contained HM markers such as *Tmem119* and *Cst3*. Thus, despite technical differences in each dataset, harmonic representations of disease-associated regions and interactions revealed the same underlying gene-level patterns – all in a fully unsupervised manner.

**Figure 9.**
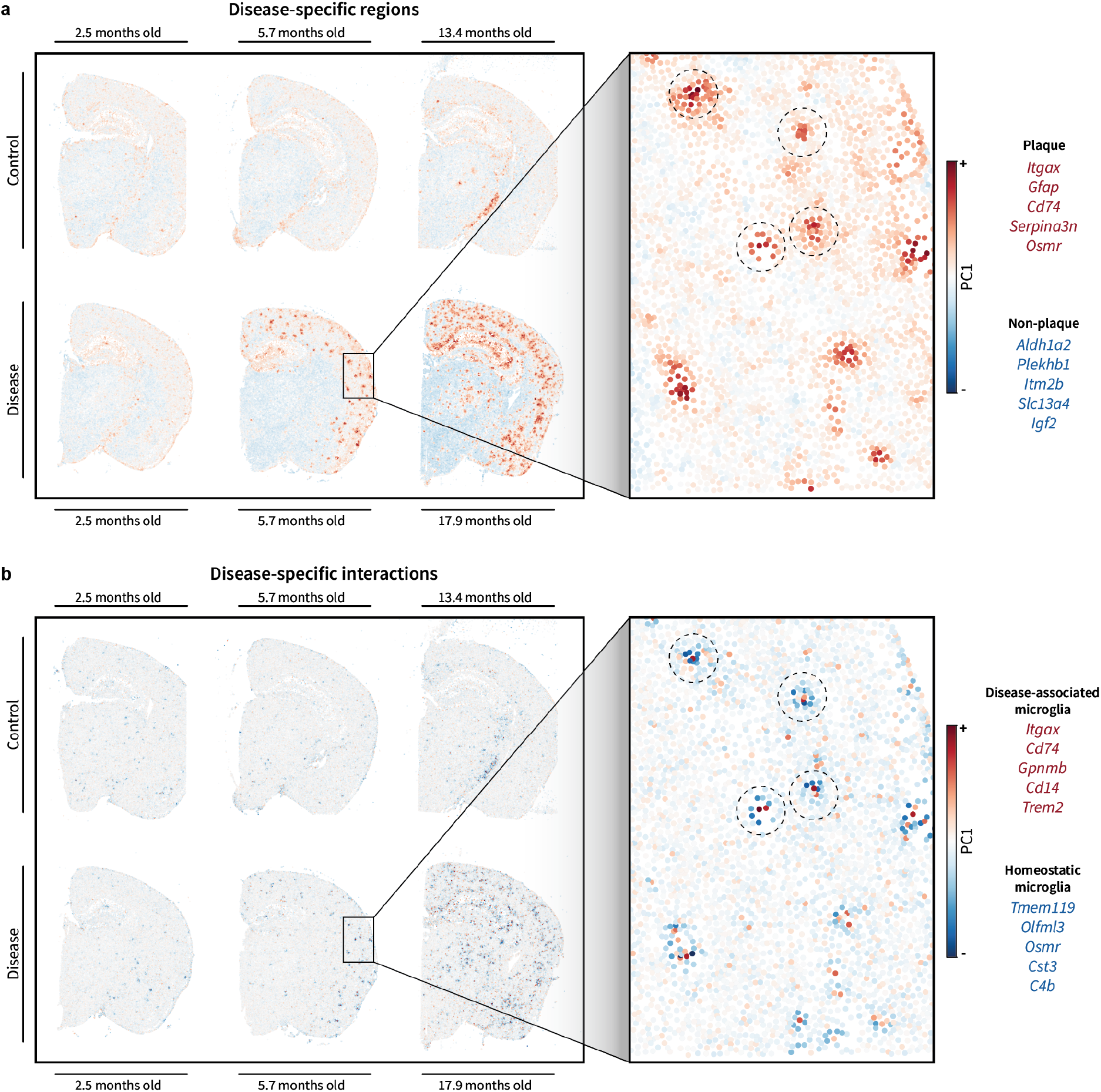
**Unsupervised identification of plaque-associated regions and interactions** in Xenium Alzheimer’s mouse model dataset. Data contains 347 genes measured across 350,219 cells total. **a**,**b)** Same as Fig. 8 but using Xenium data.

## Discussion

In this work, we have introduced representations of regions, interactions, and boundaries based on transcriptional harmonics over tissue graphs. The resulting equations revealed a duality in which regions and interactions are complementary modifications of spatial covariance, each with unique strengths and weaknesses. We derived these results from first principles and demonstrated them in multiple datasets from different technologies and biological conditions. Taken together, these results suggest that harmonic representations enable efficient, interpretable, and fully unsupervised identification of regions and interactions in spatial transcriptomics data.

We anticipate that this framework will readily extend to different types of data. For instance, despite being relatively low-plex, spatial proteomics data can also be represented as molecular signals over a spatial graph domain. Such data may also provide more accurate representations of interactions given direct measurement of the proteins that form them. Additionally, this framework could likely be generalized to the subcellular level of spatial transcriptomics data by connecting individual mRNA reads into a spatial graph domain with signals given by one-hot encodings of gene identity. Meanwhile, this work could also be extended conceptually. For example, we expect that interactions involving multiple cells could be represented via a generalization of this framework to hypergraphs. Finally, we hope that the provided interpretations might influence the choice of graph neural network architectures used in spatial transcriptomics. Just as the graph neural network literature has recently shifted focus from *homo*phily (i.e. low-frequency features) to *hetero*phily (i.e. high-frequency features) (Luan et al., 2022; Chen et al., 2024; Ma et al., 2022), perhaps spatial transcriptomics, too, might shift toward heterophilous architectures capable of representing interactions.

Altogether, we hope this work serves as a conceptual framework within which current methods can be understood as well as a quantitative foundation upon which future methods can be built.

## Acknowledgements

We thank our friends and colleagues Yanze Wang, Seth Furniss, Shuchen Luo, Ajay Nadig, Dániel Barabási, Tushar Kamath, and Ben Harris for helpful discussions and feedback on the manuscript.

## A. Supplementary Figures

**Figure A1.**
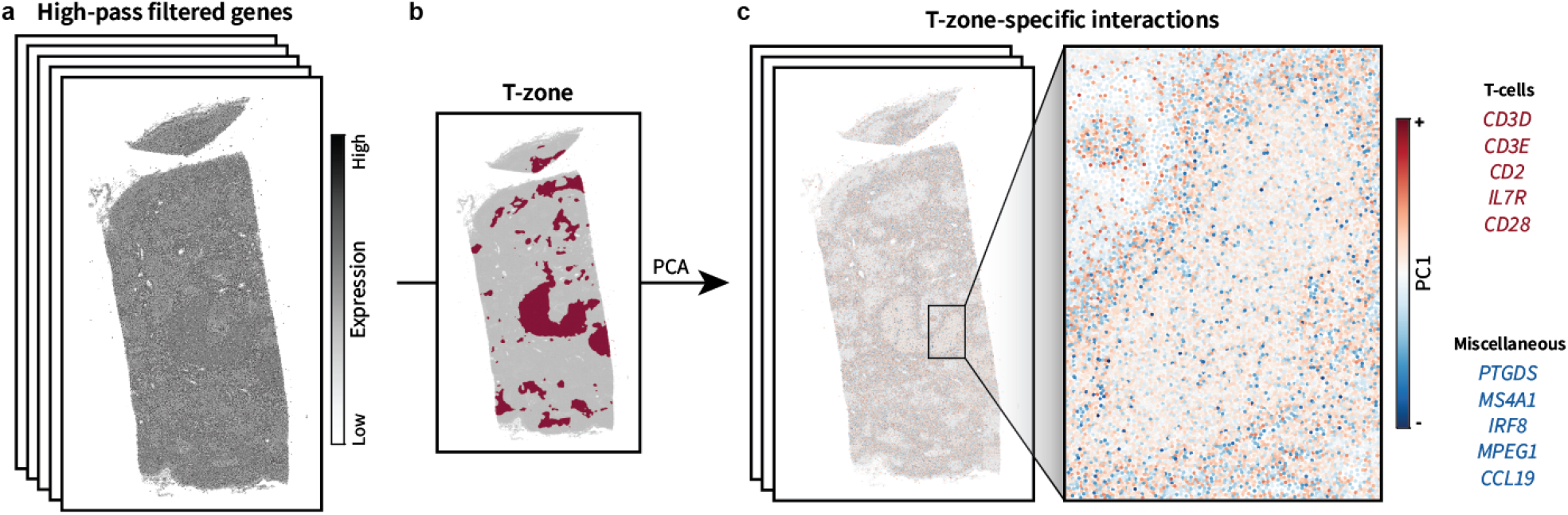
PCA applied to high-pass filtered gene expression signals in the T-zone only. **a-c)** Same as Fig. 4 but using standard PCA instead of generalized PCA.

**Figure A2.**
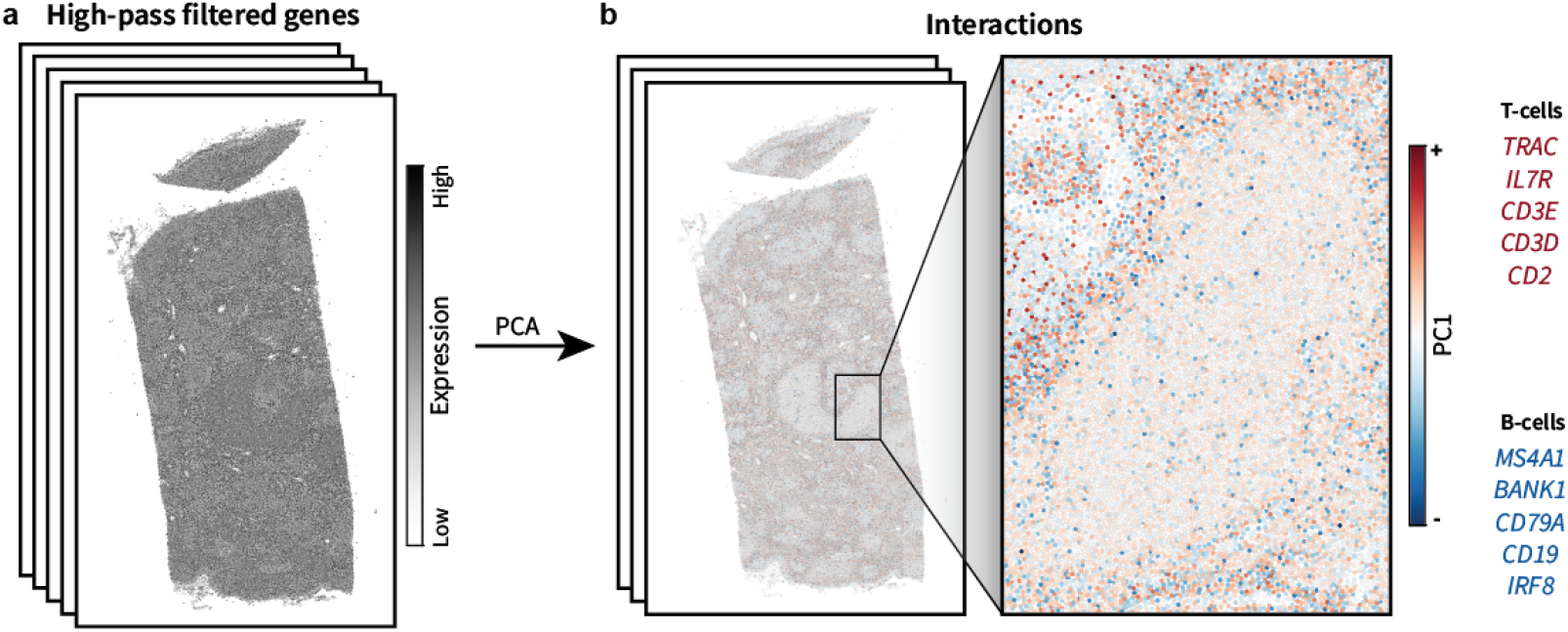
PCA applied to high-pass filtered gene expression signals. **a**,**b)** Same as Fig. 4 but using standard PCA instead of generalized PCA and without isolating the T-zone.

**Figure A3.**
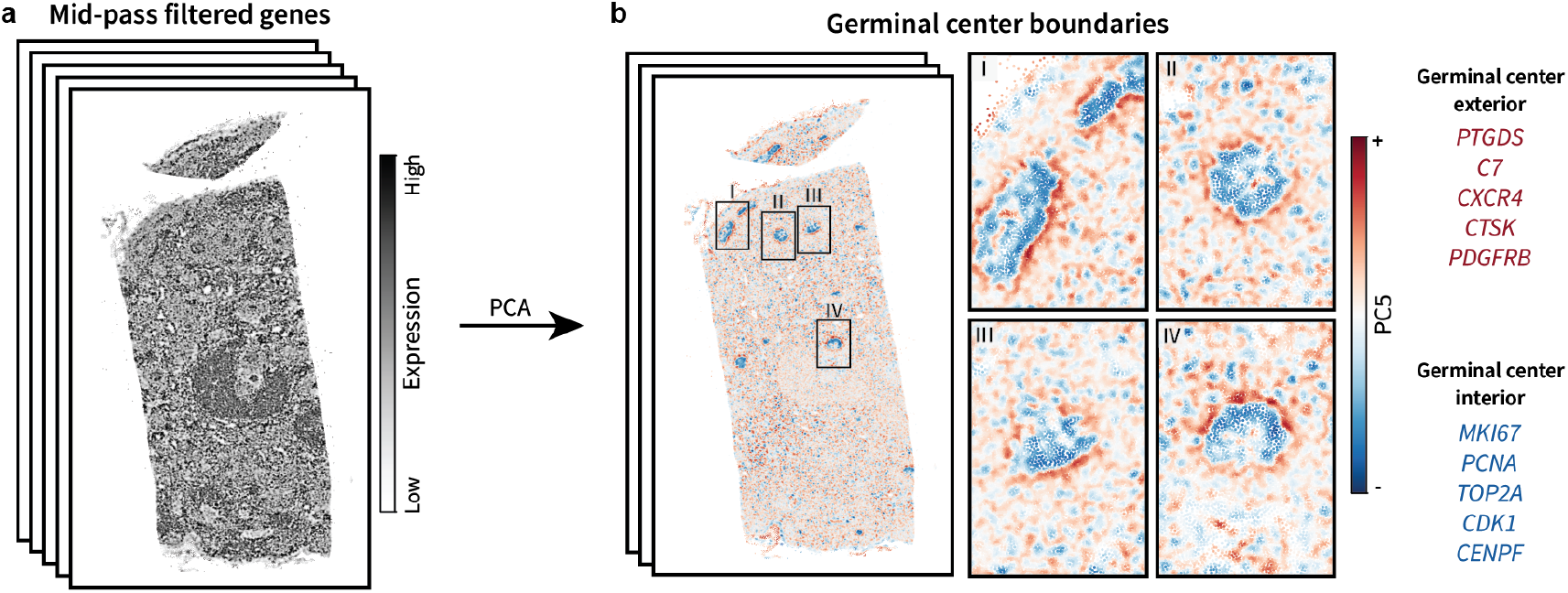
Germinal center boundaries in the human lymph node. **a)** Same as Fig. 5a. **b)** Same as Fig. 5b but showing PC4 with four different zoom-in perspectives.

**Figure A4.**
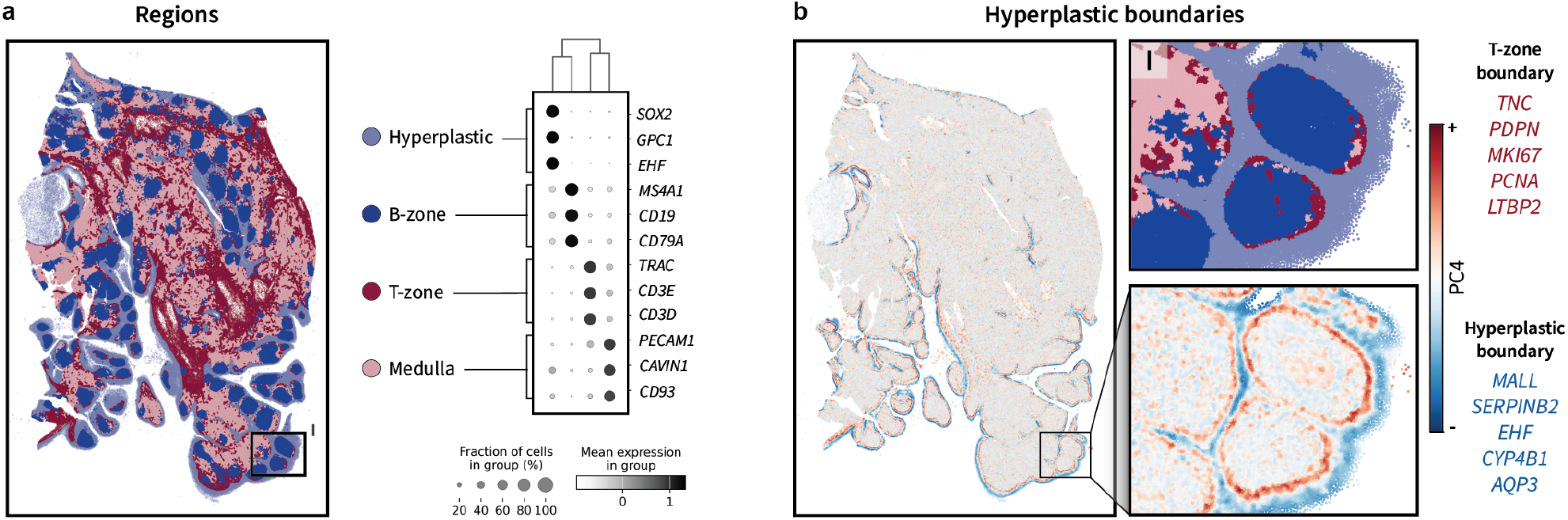
Hyperplastic boundaries in the human tonsil. **a)** Same as Fig. 7a but with an additional zoom-in for reference. **b)** Same as Fig. 7c but showing PC4 along with an additional region zoom-in above. Note that while the boundary appears to surround follicles (B-zones), it only strictly appears next to T-zones.

**Figure A5.**
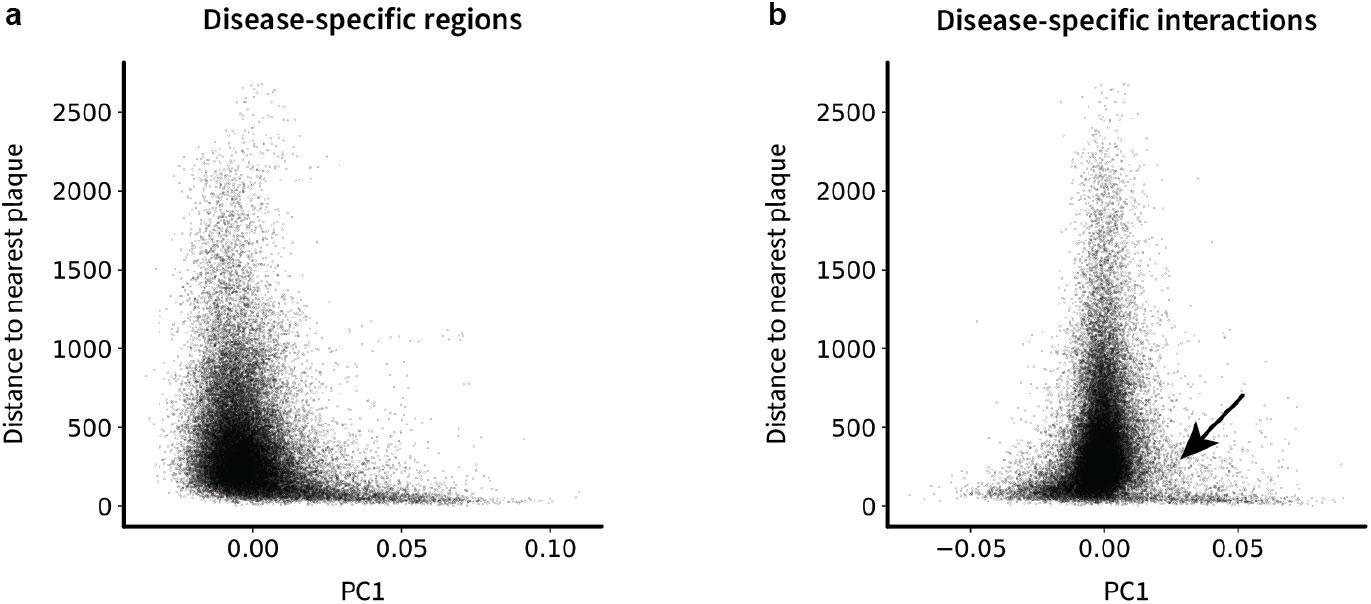
Quantification of region and interaction distances to plaques. **a)** Scatter plot showing each cell’s distance to the nearest plaque compared to its value along PC1 obtained from disease-specific region analysis (e.g. Fig. 8a). **b)** Same as a) but showing PC1 obtained from disease-specific interaction analysis (e.g. Fig. 8b). Positive PC1 values represent DAMs while negative values represent HMs. Arrow indicates that the positive portion of PC1 has a lower density of cells with higher distances from plaques, whereas the negative portion of PC1 has more cells at higher distances.

## B. Mathematical details

### B.1 Indefiniteness of the spatial covariance matrix

In spatial transcriptomics, the spatial covariance matrix is typically defined as

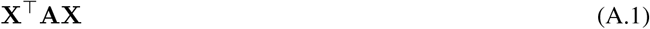

(Chen, 2015; Miller et al., 2021; Palla et al., 2022b; Russell et al., 2023). Note that the adjacency matrix, **A** (defined in Section 1.1), is indefinite. In other words, it has a mixture of positive and negative eigenvalues. To see this, first consider that it has a zero diagonal. Thus, its trace is zero. The trace of a matrix is also equal to the sum of its eigenvalues. Thus, the sum of the eigenvalues of **A** is zero. One might consider the possibility that all the eigenvalues are zero. However, that is not a possibility here because **A** contains at least some nonzero entries, i.e. at least some connections between cells. Thus, the eigenvalues of **A** must be a mixture of positive and negative in order to cancel out to zero, and **A** is indefinite. Note that the same holds true for spatial covariance, **X**^⊤^**AX**.

In the context of PCA, indefiniteness presents two problems. First, the eigenvalues of the spatial covariance matrix correspond to the “variance explained”. If the eigenvalues are negative, this prevents interpretation in terms of variance explained. Second, PCA is typically implemented via singular value decomposition (SVD) of the original data matrix. In this case, the original data matrix would be

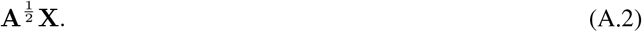

One can verify this by seeing that matrix multiplication with itself indeed produces the desired spatial covariance matrix. However, the matrix 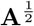 can be expressed in terms of diagonalization as

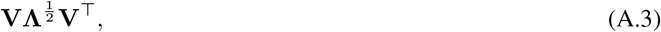

where **V** is the eigenbasis of the adjacency matrix, and **Λ** is the diagonal matrix of its eigenvalues. As **A** is indefinite, some of its eigenvalues are negative, and we end up taking the square root of negative numbers, producing complex values. This leads to downstream complications regarding computation and interpretation. The same issue arises independently of SVD in that one must calculate this original data matrix in order to project it onto the resulting eigenbasis to calculate PC embeddings.

Altogether, it is for the above reasons that we claim PCA cannot be performed using the spatial covariance matrix.

### B.2 Frequency summation as a quadratic form

Eq. (1) can be converted into a quadratic form via an expansion and then a contraction of the summation. First, factor the **v** terms to get

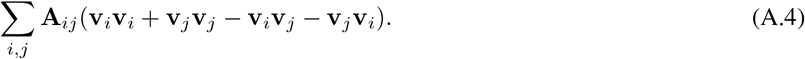

The positive and negative terms can then be combined, respectively, yielding

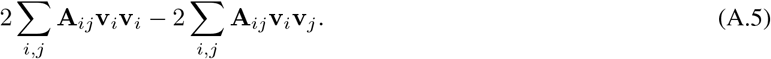

Note that, for a fixed row *i*, the left-hand term corresponds to summing **v**_*i*_**v**_*i*_ together *d*_*i*_ times, where *d*_*i*_ is the sum of the *i*th row of **A**. Thus, one could equivalently express the left-hand term as

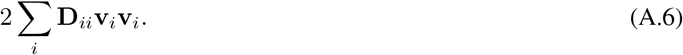

Now both the left and right terms themselves correspond to quadratic forms, yielding

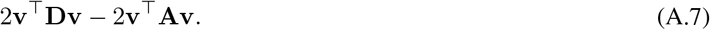

Further simplifying, we have

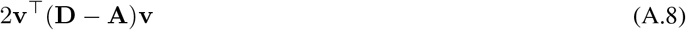

and further

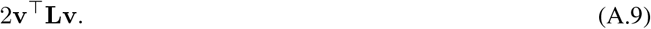

The factor of two is conventionally omitted, yielding the final expression

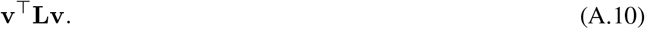

### B.3 Interaction summation as a quadratic form

Conversion of Eq. (6) into a quadratic form can be performed using the same steps as in Section B.2. First, factor the x and **y** terms to get

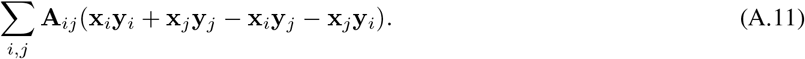

The positive and negative terms can then be combined, respectively, yielding

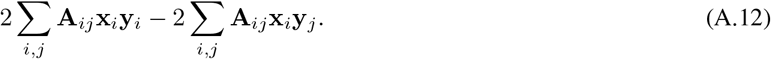

This simplifies to

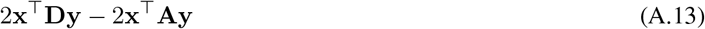

and further

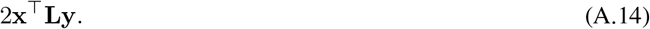

Again omitting the factor of two by convention, we have the final expression

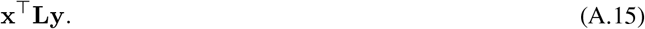

### B.4 Additional interpretation of generalized eigendecomposition

An additional interpretation of a generalized eigenvalue problem can be gained by expanding it into its corresponding Rayleigh quotient. A standard eigendecomposition, using the T-zone-specific covariance matrix 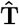, is given by

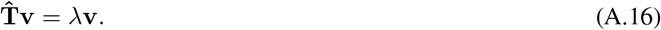

Multiplying each side by **v**^⊤^, we have

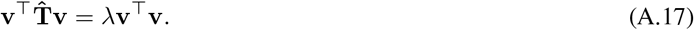

Solving for *λ* gives the Rayleigh quotient

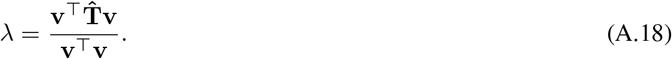

This eigenvalue corresponds to the “amount of variance explained”. Thus, the eigenvalue corresponding to the maximum variance explained, i.e. PC1, is

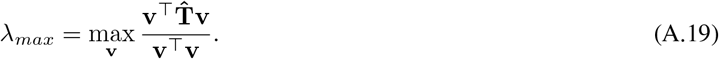

This can be interpreted as finding the “axis” **v** that captures as much information in 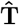 as possible. Note that the denominator simply serves to normalize the expression such that vector magnitude is not considered.

So far, this describes standard PCA. However, this approach can be generalized by considering a second matrix, in our case **Ŝ**. As explained in Section 3.2, we seek to *divide out* patterns outside the T-zone (given by **Ŝ**) in order to find patterns present uniquely present within the T-zone (given by 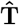). We can incorporate this intuition directly into the Rayleigh quotient above by calculating

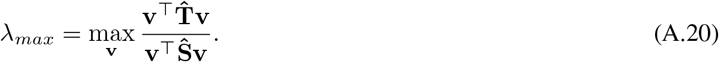

In this form, one can see that we are explicitly dividing out **Ŝ**. As a result, any patterns captured by **v** that are also present in **Ŝ** result in a larger denominator and thus a smaller *λ*. Thus, in order to maximize *λ*, the corresponding **v** must strongly capture patterns that are present within 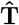 but not **Ŝ**. Finally, reversing the expansion performed above, we can convert this Rayleigh quotient into the generalized eigenvalue problem

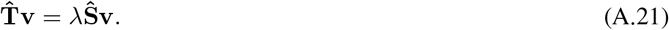

### B.5 L and Γ share the same eigenvectors with reversed eigenvalues

In Section 5.1, we introduced the matrix **Γ** = *λ*_*max*_**I** − **L**. This matrix has the same eigenvectors as the Laplacian but with effectively reversed assignment of eigenvalues. To see this, consider two facts. First, negation of a matrix maintains the same eigenvectors while simply negating the eigenvalues:

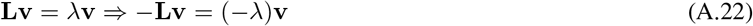

Second, addition of a multiple of the identity matrix simply shifts the eigenvalues by that multiple:

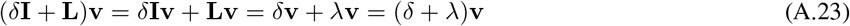

Because **Γ** is equal to negating the Laplacian and shifting it up by a multiple of the identity matrix, its eigenvectors are the same as those of the Laplacian. Its eigenvalues, however, are negated and shifted up by *λ*_*max*_, thus effectively reversing their assignment to each eigenvector.

### B.6 Region quadratic form as a summation

Conversion of Eq. (16) into a the summation given by Eq. (17) follows the same steps as in Sections B.2 and B.3 but in the opposite order. We begin with

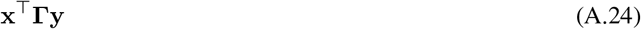

and add the factor of two that is typically omitted for convention:

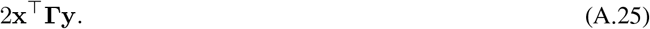

Recall that

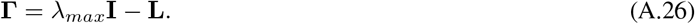

Given the assumption that *λ*_*max*_ = 2, we have

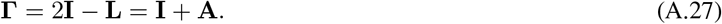

Thus, eq. (A.25) can be expanded into

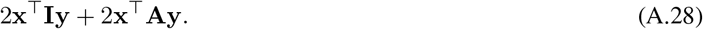

Writing each quadratic form as a separate sum, we have

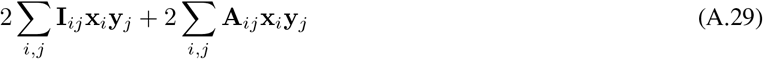

and, equally,

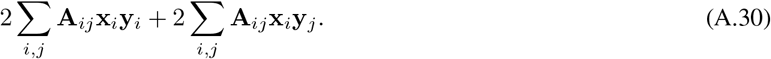

Finally, we combine each summation to get

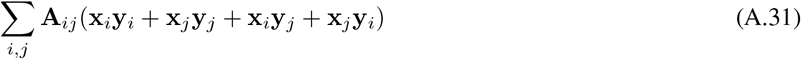

and undo the factorization to reach

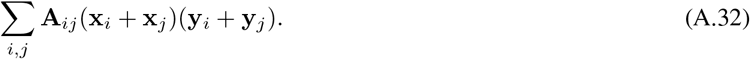

## C. Cell typing results

Here, we provide cell typing results for each dataset shown. Note that conventional cell typing corresponds to the same analyses used in the main text but without filtering. In other words, it corresponds to using the entire frequency spectrum rather than isolating lows, highs, or mids.

We performed conventional cell type clustering for each dataset by applying PCA to normalized, log-transformed, and scaled counts data followed by Leiden clustering using `sc.pp.neighbors` and `sc.tl.leiden`. The Leiden resolution was set such that cell types considered relevant to the reported interactions were identified. Cluster labels were manually assigned based on the top three marker genes for each cluster and restricted to the minimal possible interpretation sufficient to evaluate results in the main text.

**Figure A6.**
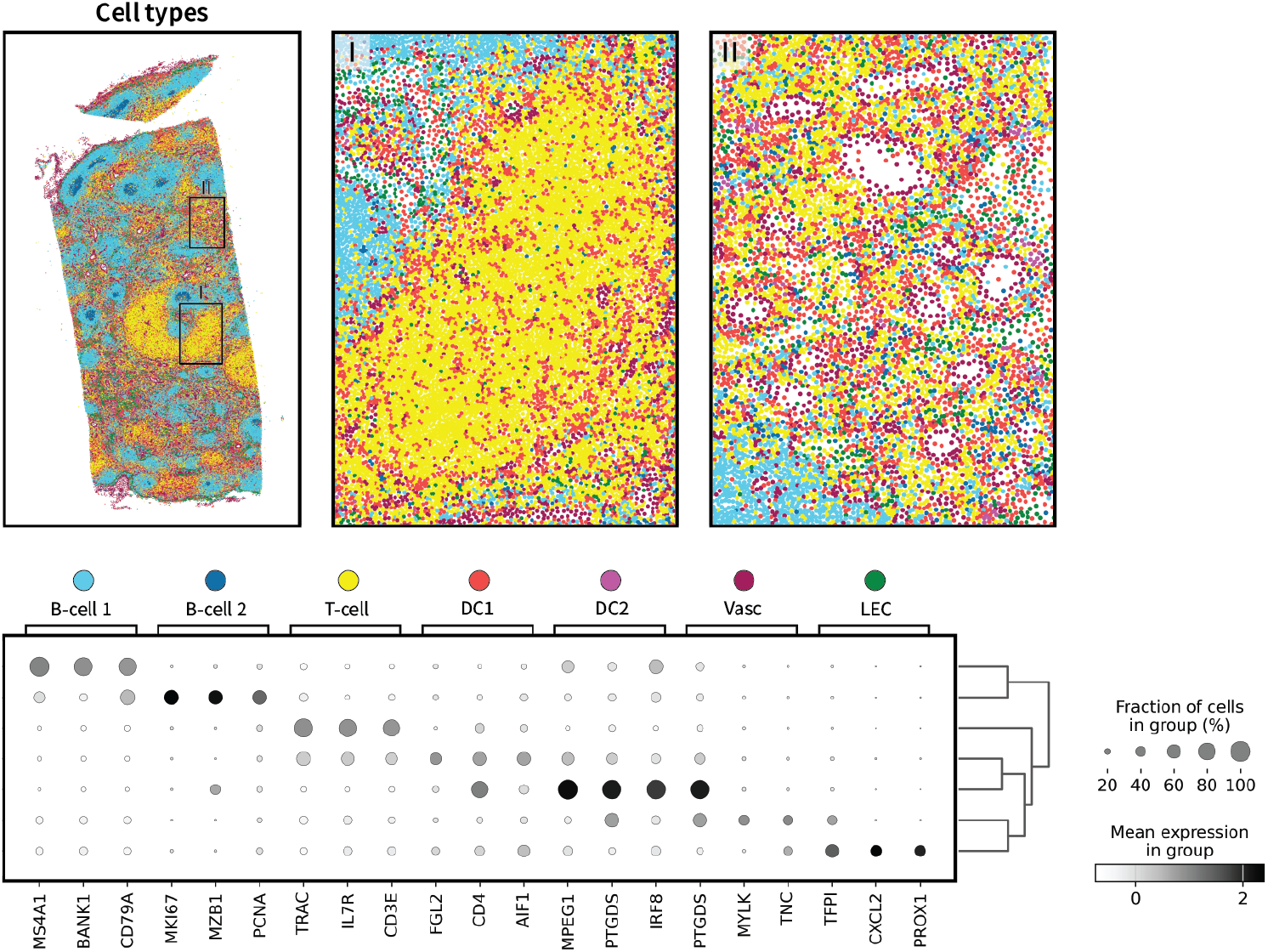
**Conventional cell type clusters** in the human lymph node dataset. DC: “Dendritic cell”, Vasc: “Vasculature”, LEC: “Lymphatic endothelial cell”.

**Figure A7.**
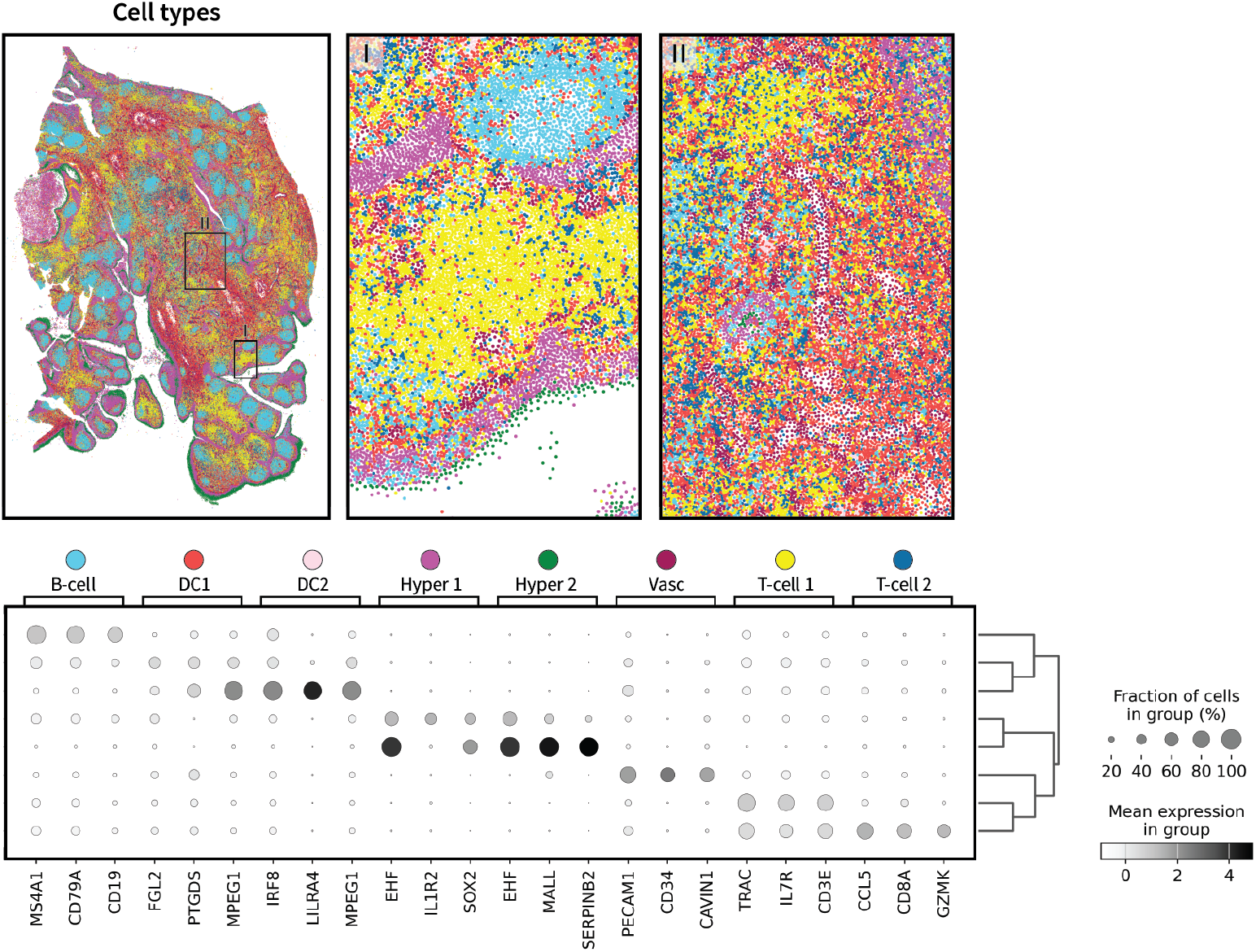
**Conventional cell type clusters** in the human tonsil dataset. DC: “Dendritic cell”, Vasc: “Vasculature”, Hyper: “Hyperplastic cell”.

**Figure A8.**
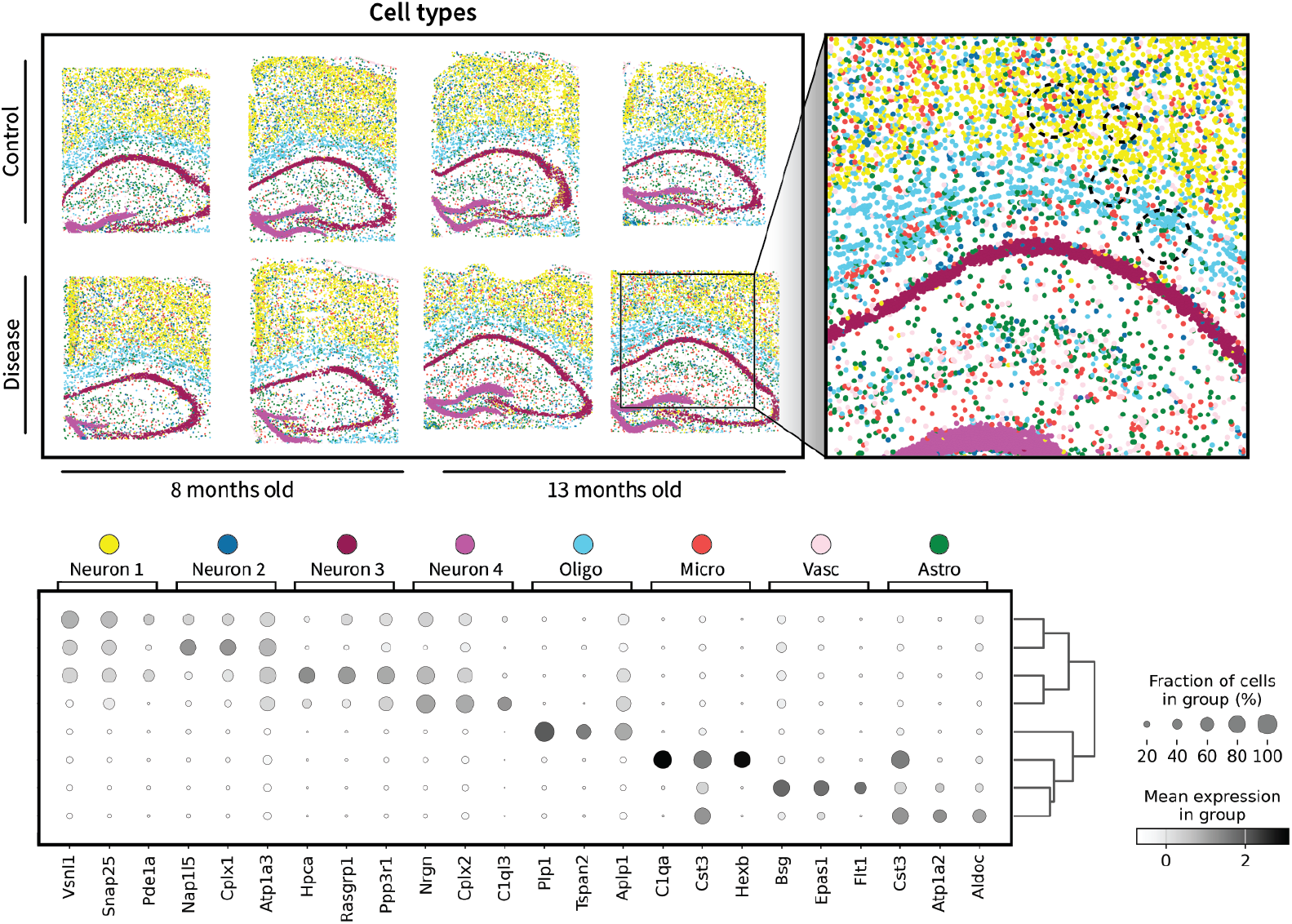
**Conventional cell type clusters** in the STARmap mouse brain dataset. Oligo: “Oligodendrocyte”, Micro: “Microglia”, Vasc: “Vasculature”, Astro: “Astrocyte”.

**Figure A9.**
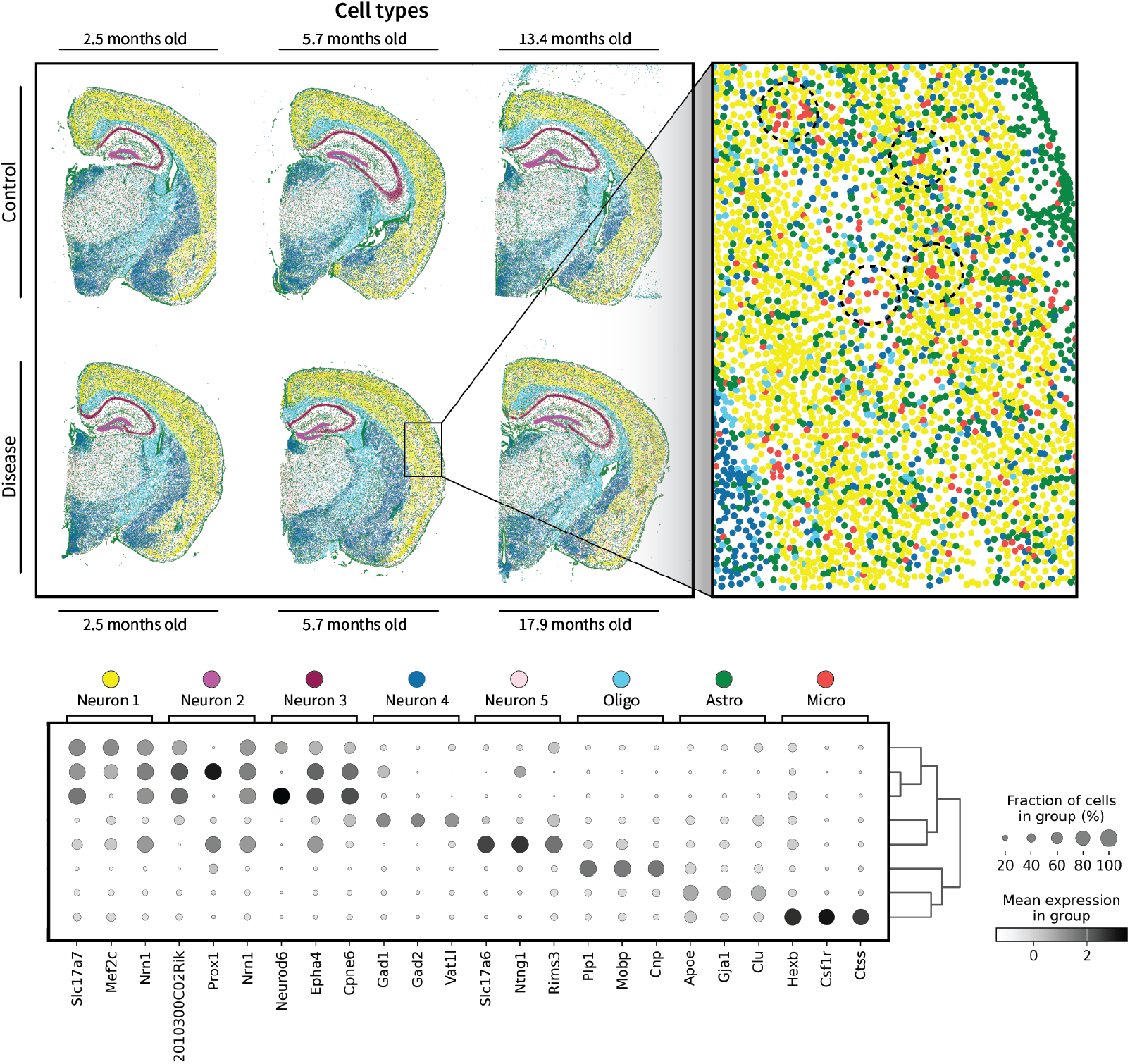
**Conventional cell type clusters** in the Xenium mouse brain dataset. Oligo: “Oligodendrocyte”, Micro: “Microglia”, Astro: “Astrocyte”.

## D. Connections to image processing and graph neural networks

Here, we mention connections between the proposed harmonic representations and related themes in other fields. In particular, we make connections to image processing and graph neural networks, as they are quantitatively similar and may provide further intuition as well as inspiration for the development of future methods.

### D.1 Regions

#### Image processing

The diffusion kernel corresponds to a Gaussian blur in image processing. Gaussian blurring is often used to reduce noise in images by making nearby pixel values more similar to one another. In other words, it is a low-pass filter. Thus, the application of the diffusion kernel above can be thought of as a Gaussian blur over a graph. It is likely this analogy that leads to the notion that low-pass filtering reduces the amount of noise in spatial transcriptomics data (Long et al., 2023; Xu et al., 2024). However, we have argued in the main text that high-frequency patterns in spatial transcriptomics data are not simply noise.

#### Graph neural networks

Graph neural networks (GNNs) can be understood as iterative low-pass filters over a graph (Kipf & Welling, 2017; Wu et al., 2019). Additionally, they can *learn* relevant kernels based on training data. An important assumption underlying conventional GNNs is that nearby nodes have similar features, i.e. “homophily”. This is the inductive bias that leads to the low-pass nature of GNNs.

### D.2 Interactions

#### Image processing

The Laplacian filter (i.e. performing **L**x on a signal x) is a cornerstone of image processing, as it enables edge detection in images. Indeed, the interactions described in the main text could instead be understood as opposing molecular edges between neighboring cells. In an image, this could be pictured as an edge going from high to low in the red channel while the blue channel forms an edge going from low to high.

#### Graph neural networks

Recent work on GNNs has shifted toward the assumption of *hetero*phily (i.e. “opposites attract”), leading to models that act as *high*-pass filters (Luan et al., 2022; Chen et al., 2024; Ma et al., 2022). While these networks may be suitable for detecting interactions in spatial transcriptomics data, to our knowledge, they have yet to be applied.

### D.3 Boundaries

#### Image processing

Just as high– and low-pass filtering were combined to define region boundaries, so can Laplacian and Gaussian filters be combined to form the Laplacian of Gaussian (LoG) filter in image processing. This conventional filter enables one to reduce noise using a Gaussian filter before identifying edges via a Laplacian filter. This can also be understood as identifying edges on a given length scale determined by the Gaussian filter.

#### Graph neural networks

High– and low-pass filtering can also be combined to create GNNs that address both homophily and heterophily (Chen et al., 2024). Examples of such approaches include reaction-diffusion-based networks, in which high-pass filtering corresponds to reaction of a given feature with itself, while low-pass filtering corresponds to diffusion (Choi et al., 2023). We find these architectures particularly interesting due to their inherent modeling of the underlying dynamics of the features over the graph. This could open the door to rational design of filter kernels (or, equivalently, inductive biases for GNNs) using experimentally-derived biophysical principles.

## Notes

### Competing Interest Statement

The authors have declared no competing interest.

